# Dorsoventral dissociation of Hox gene expression underpins the diversification of molluscs

**DOI:** 10.1101/603092

**Authors:** Pin Huan, Qian Wang, Sujian Tan, Baozhong Liu

## Abstract

Unlike the Hox genes in arthropods and vertebrates, those in molluscs show diverse expression patterns and, with some exceptions, have generally been described as lacking the canonical staggered pattern along the anterior-posterior (AP) axis. This difference is unexpected given that almost all molluscs share highly conserved early development. Here, we show that molluscan Hox expression can undergo dynamic changes, which may explain why previous research observed different expression patterns. Moreover, we reveal that a key character of molluscan Hox expression is that the dorsal and ventral expression is dissociated. We then deduce a generalized molluscan Hox expression model, including conserved staggered Hox expression in the neuroectoderm on the ventral side and lineage-specific dorsal expression that strongly correlates with shell formation. This generalized model clarifies a long-standing debate over whether molluscs possess staggered Hox expression and it can be used to explain the diversification of molluscs. In this scenario, the dorsoventral dissociation of Hox expression allows lineage-specific dorsal and ventral patterning in different clades, which may have permitted the evolution of diverse body plans in different molluscan clades.

---

A conserved role of Hox genes in body patterning across bilaterian animals has been extensively discussed^1–3^ and even extended to cnidarians to some extent^4,5^. However, conservation has limits, and there is considerable variation between animal lineages in the composition and expression patterns of Hox genes^6–12^. These differences are particularly evident in Spiralia, which together with Ecdysozoa and Deuterostomia, forms the three major clades of Bilateria. With the exception of Annelida, which shows clear canonical staggered Hox expression along the anterior-posterior (AP) axis (similar to arthropods and vertebrates)^13–15^, staggered Hox expression has been observed less frequently in spiralians than in other bilaterian clades^10,11,16,17^.

Mollusca is the most species-rich phylum of Spiralia and comprises seven or eight “class”-grade clades^18–20^ (Fig. 1a). Hox expression in molluscs shows diverse patterns. Although staggered expression has been reported, it is observed in a variety of tissues in different clades^21–24^. The observation of a staggered Hox pattern in phylogenetically distant molluscan clades suggests that staggered Hox expression is widespread in molluscs. However, it remains unclear why such expression has been observed in some species (polyplacophorans, scaphopods and bivalves)^21–24^ but not others (e.g., in gastropods^25–27^ and cephalopods^28^, although evidence has recently suggested staggered expression in the two clades^23^). In addition, even without seeking a common staggered pattern, molluscan Hox expression is still too diverse to conclude a general model. Although correlations between Hox expression and the larval shell field, foot and nervous system are frequently observed^21–28^, the Hox expression in these particular organs shows few common patterns except that the expression in the nervous system in late-stage larvae seems to be common in different clades^22,23,25,28^.

**Fig. 1 a.**
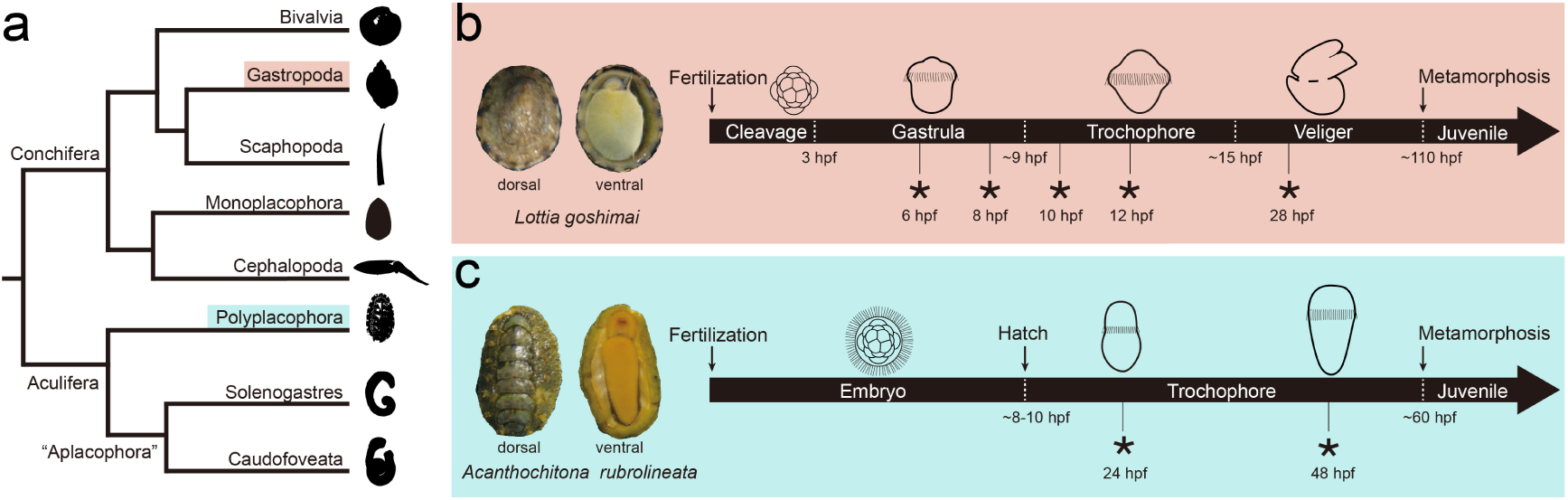
A phylogenetic tree of Mollusca (based on a previous article^18^). The phylogenetic positions of the two species used in this study (the gastropod and polyplacophoran) are highlighted. **b and c.** The gastropod *Lottia goshimai* and the polyplacophoran *Acanthochitona rubrolineata*. The morphology of adults and the timing of development (length not to scale) are shown. Asterisks indicate the time points at which samples were collected for gene expression analysis in the present study.

One possible explanation for the diverse Hox expression in molluscs is that the Hox genes may have been recruited to regulate the development of lineage-specific structures (lineage novelties). This recruitment might, for example, underpin the roles of Hox genes in the development and evolution of the brachial crown in cephalopods^28^. However, despite the contribution of lineage-specific expression, the great diversity of molluscan Hox expression is still unexpected given that all molluscs share highly similar early development, including spiral cleavage and highly conserved primary larvae (trochophore) (the conchiferan molluscs also share late larvae, or veligers)^29–32^. We propose an additional explanation for the diverse molluscan Hox expression patterns: there could be some “cryptic” similarities in Hox expression that remain unrevealed. In this scenario, Hox expression may undergo dynamic changes during development, and different molluscan lineages may share common expression patterns at particular stages. Dynamic Hox expression in molluscs is supported by the fact that distinct Hox expression patterns can be observed at different developmental stages in a single species^23,25^. If correct, such dynamic expression may explain why staggered expression has been observed in some species but not in others. Indeed, previous studies on molluscan Hox expression focused on very different developmental stages^21–28^, which may have contributed to the poor consistency of the results.

To test our hypothesis, it is necessary to investigate Hox gene expression at multiple developmental stages in individual molluscan species and then compare expression among different clades. Here, we performed a comprehensive investigation of various developmental stages of a gastropod mollusc (the limpet *Lottia goshimai*) (Fig. 1b). We paid special attention to a gastropod species because previous studies on gastropod Hox expression revealed the most diverse results and a general lack of staggered expression^25–27^. For comparison, we included a polyplacophoran, the chiton *Acanthochitona rubrolineata* (Fig. 1c), which belongs to the aculiferan molluscs and is phylogenetically distant from *L. goshimai* (Fig. 1a).

## Results

### The molluscan Hox: general remarks

We identified 11 and 10 Hox genes from the developmental transcriptomes of *L. goshimai* and *Ac. rubrolineata*, respectively (Fig. 4c and supplemental figures S13-S14). The two species both possessed complete anterior (*hoxs1-3*) and central (*hoxs4-5, lox5*, *antp*, *lox4* and *lox2*) classes of Hox genes. For the posterior class, two genes were identified in *L. goshimai* (*post2* and *post1*), whereas only *post2* was observed in *Ac. rubrolineata.* No *post1* gene was retrieved when we further searched against an adult transcriptome. Given that *post1* is also not found in the chiton *Acanthochitona crinita*^22^, this gene may have been lost in this lineage.

In *L. goshimai*, we sampled five developmental stages (two gastrula stages, two trochophore stages and one veliger stage) in short time intervals (as short as every two hours) and confirmed that its Hox expression changed rapidly (see Fig. 2 for examples). These changes correlated with early developmental events, including gastrulation as well as shell and foot development (supplemental figures S1-S4). In the late larva (28 hours post fertilization (hpf) veliger), the expression of most Hox genes was detected in the neural tissues of the foot and internal organs (supplemental figure S5). In *Ac. rubrolineata*, we sampled two larval stages and detected minor changes in Hox expression. Although other developmental stages (e.g., the embryos and very early larvae) may exhibit different Hox expression, we believe that the two stages are sufficiently representative because they showed Hox expression patterns similar to those of *L. goshimai* (see details below). In the late larva (48 hpf, comparable to the veliger larva of *L. goshimai*), Hox expression was readily detected in the shell field in addition to high Hox expression in the neural tissues in the foot. Other noticeable results included asymmetrical left-right expression and mesodermal expression of some *L. goshimai* Hox genes. Nevertheless, because we focused on an inter-species comparison in the present study, we will not describe the details of each Hox expression pattern here; instead, we provide this information in supplemental figures S1-S7.

**Fig. 2.**
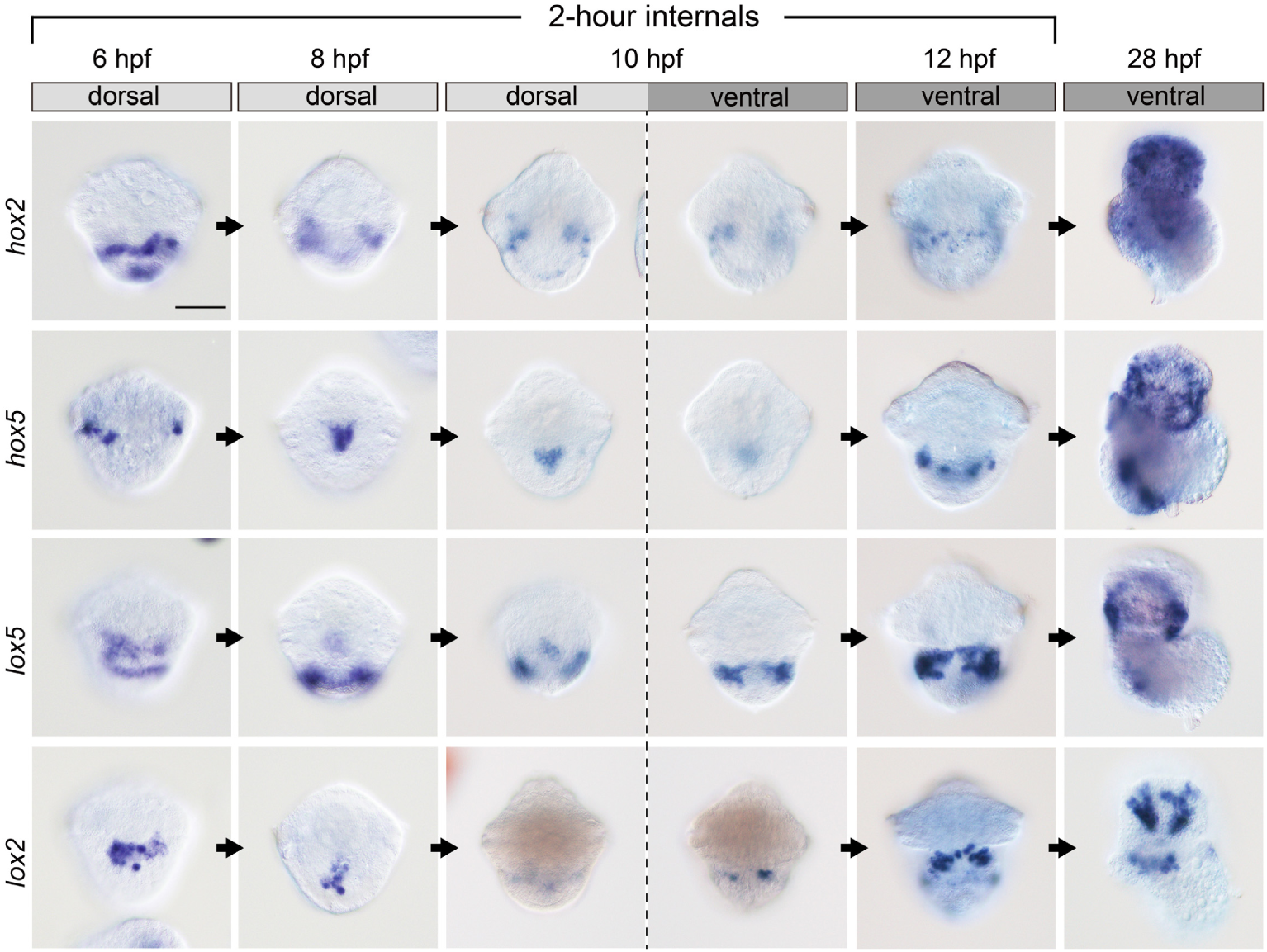
Hox expression in *L. goshimai* changes rapidly in early development. Here, four genes are presented as examples, and information on other genes is provided in supplemental figures S1-S5. A dorsal view is shown for 6-8 hpf, and a ventral view is shown for 12-28 hpf, as these are the views showing the greatest Hox expression. Both dorsal and ventral views are shown for 10 hpf. The bar represents 50 μm.

### The key characteristic of molluscan Hox expression: dorsoventral dissociation

After analysing the complex Hox expression at multiple developmental stages, we concluded that a key characteristic of molluscan Hox expression is the dissociation of dorsal and ventral expression (see examples in Fig. 3). At a given time point, a Hox gene can be expressed solely in dorsal or ventral tissues (e.g., *L. goshimai hox5* and *lox4* at 10 hpf, see Fig. 3e, h) or in both tissues while showing distinct expression patterns (e.g., *L. goshimai hox4* at 10 hpf and most *Ac. rubrolineata* Hox genes at 24 hpf, see Fig. 3d, i-u). This spatial dissociation is crucial for recognizing both conserved and lineage-specific patterns of molluscan Hox expression. As we will describe in detail below (Figs. 4 and 5), in both molluscan species, ventral Hox expression may contribute to neuroectoderm/foot development, and dorsal expression correlates with shell formation.

**Fig. 3.**
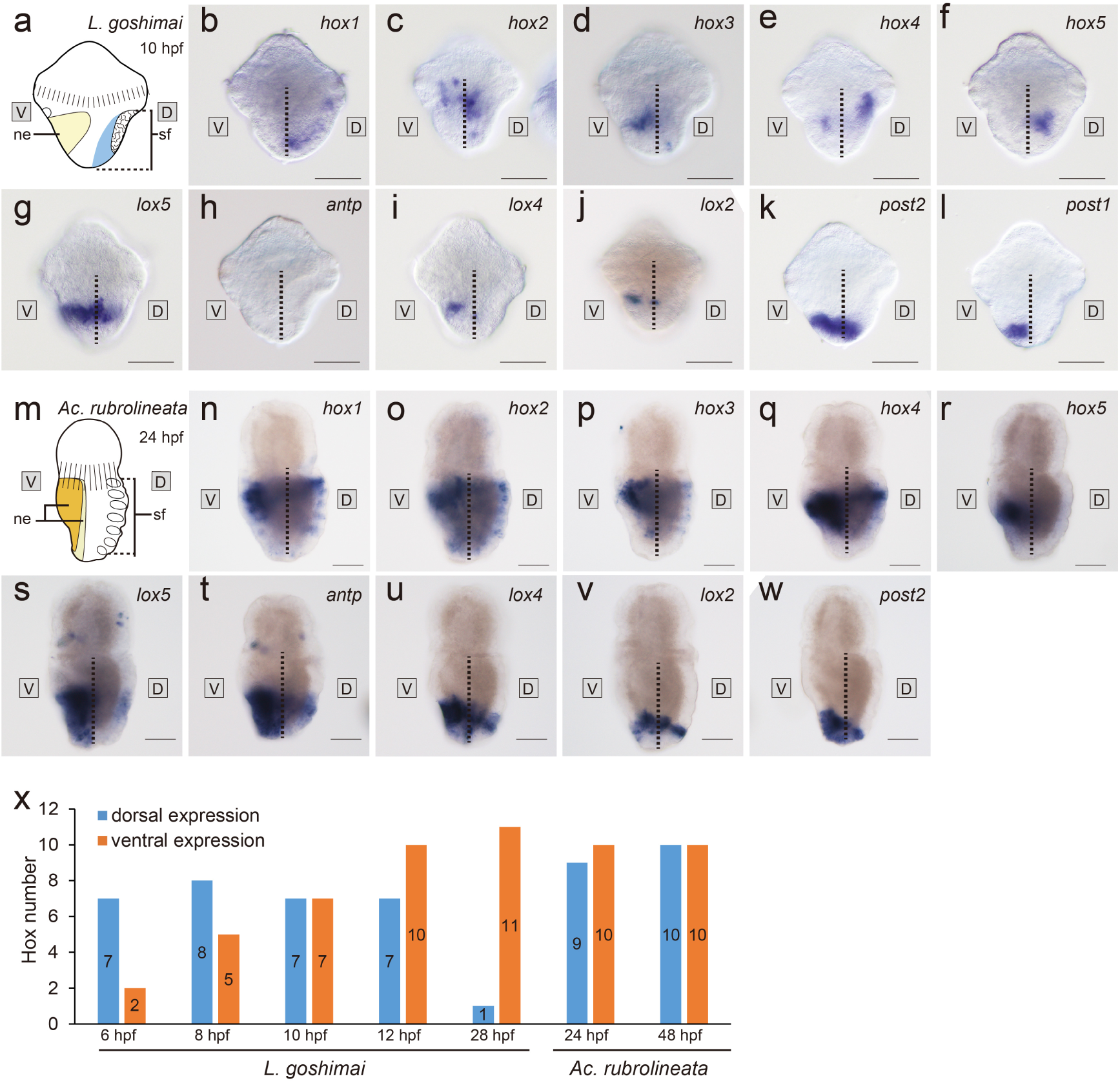
Dorsoventral dissociation of molluscan Hox expression. All panels except **x** are lateral views. Spatial dissociation is observed in both species (**b-l**: *L. goshimai*; **n-w**: *Ac. rubrolineata*), while temporal dissociation is observed in *L. goshimai* but not obvious in *Ac. rubrolineata* (**x**). The dashed lines separate the dorsal (D) and ventral (V) parts in panels **b-l** and **n-w**. Panels **a** and **m** show schematics of the trochophore larvae of the two species (lateral view), respectively. ne, neuroectoderm; sf, shell field. Bars represent 50 μm. Here, two representative stages (one for each species) are provided as examples, and all data are provided in supplemental figures S1-S7.

**Fig. 4.**
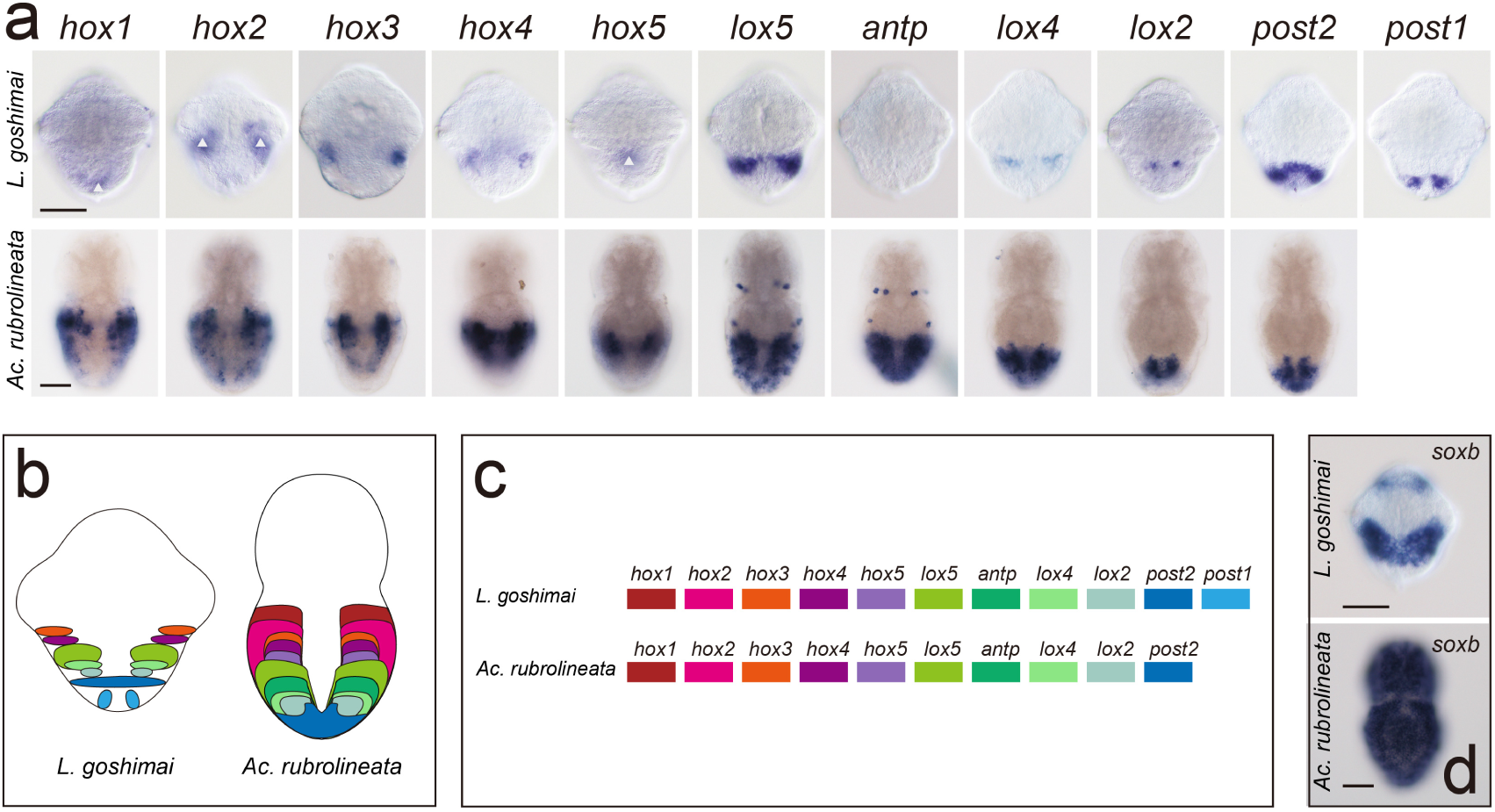
Conserved staggered Hox expression along the AP axis of the neuroectoderm. The larvae are at the same stage as in Fig. 3a-w, and all panels are ventral views. Panel **a** shows the Hox expression in the trochophores of *L. goshimai* and *Ac. rubrolineata*. Schematic diagrams are presented in **b** that show the staggered Hox expression along the AP axis according to a presumptive Hox cluster (**c**). Note that expression of *hox1*, *hox2* and *hox5* of *L. goshimai* (white triangles) is not neuroectodermal and expression of *antp* is not detectable (see details in supplemental figure S3). Panel **d** shows the *soxb* expression in the two species, indicative of the neuroectoderm. In *L. goshimai*, *soxb* expression does not cover the most posterior ectoderm that expresses *post2* and *post1*. Bars represent 50 μm.

**Fig. 5.**
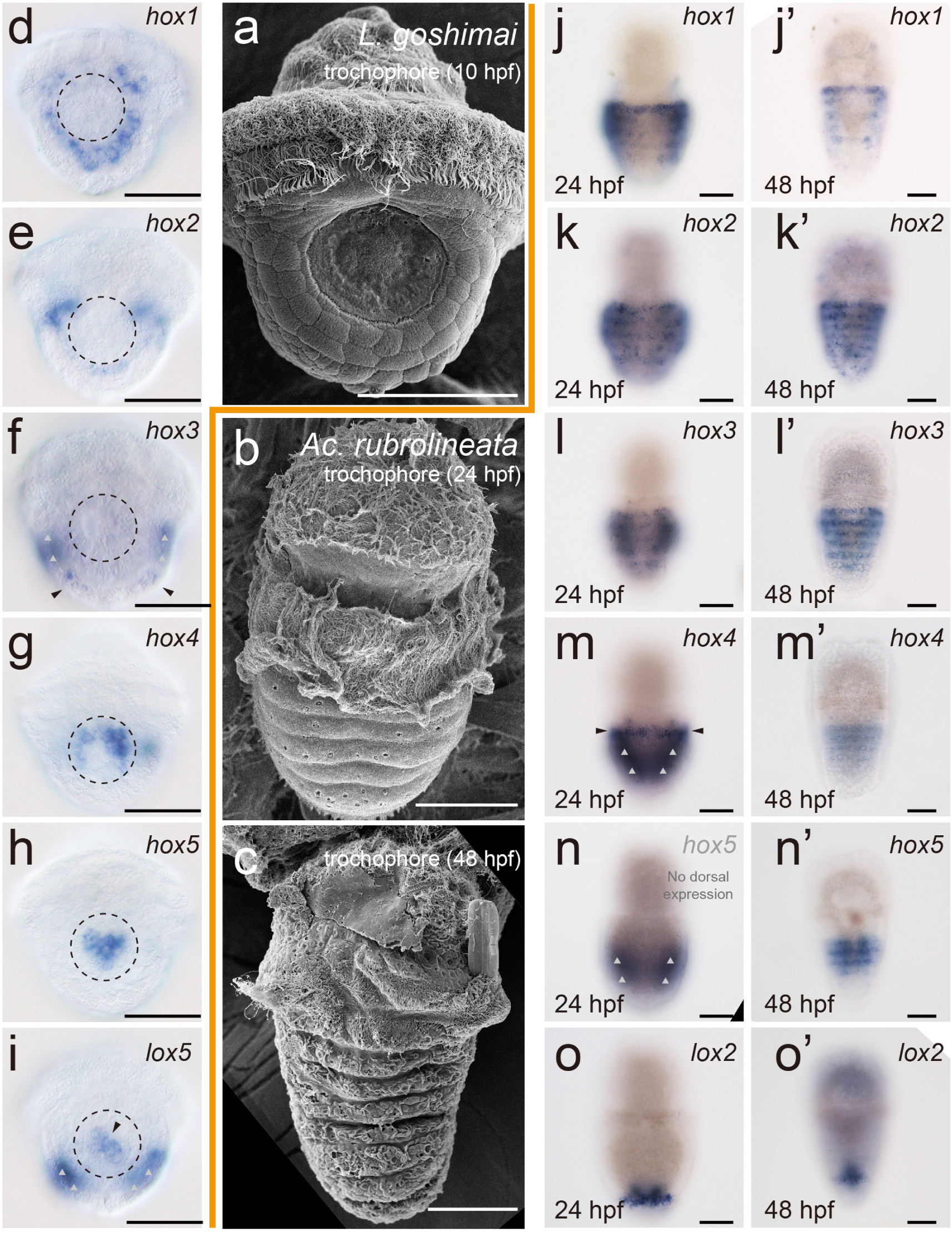
Dorsal Hox expression was correlated with the lineage-specific shell field in *L. goshimai* and *Ac. rubrolineata.* Most larvae are at the same developmental stage as in Fig. 3a-w (except **j’-o’**), and all panels are dorsal views. Panels (**a-c**) show the scanning electronic microscope images of the larvae of the two species. The round shell plate in *L. goshimai* (**a)** and the seven pseudosegments where the shell plates will form in *Ac. rubrolineata* (**b-c**) are easily distinguished. The dashed circles in **d-i** indicate the shell field. In some panels (**f**, **i**, **m** and **n**), the dorsal and ventral expression is indicated by black arrowheads and grey triangles, respectively, since the ventral expression is very intense and may interfere with the discrimination between dorsal and ventral expression. See the text and supplemental figures S1-S7 for dorsal expression of other Hox genes. Bars represent 50 μm.

In addition to spatial dissociation, Hox expression in *L. goshimai* also showed obvious temporal dissociation. As shown in Fig. 3x, the dorsal and ventral expression showed no obvious correlation in *L. goshimai*, and there was a tendency for earlier dorsal expression. Most Hox genes were expressed in dorsal tissues at early developmental stages (6-8 hpf) and were activated in ventral tissues later (after 10 hpf), although the dorsal expression was still sustained to a certain level in late stages (Fig. 3x and supplemental figures S1-S5). Two exceptions included *antp*, which showed only ventral expression (in the late larva), and *lox4*, which showed earlier ventral expression (supplemental figures S2-S5). The dorsal and ventral Hox expression in *Ac. rubrolineata* did not show temporal dissociation at the two developmental stages that were investigated (Fig. 3x and supplemental figures S6-S7).

### Ventral Hox expression: conserved staggered expression in the neuroectoderm

Ventral Hox expression showed conservation in different molluscan clades and was correlated with the development of the neuroectoderm (the majority of which later developed into the foot). In *Ac. rubrolineata*, ventral Hox expression showed a staggered pattern similar to that previously reported in its close relative (*Ac. crinita*)^21,22^. In *L. goshimai*, despite the quick changes, we recognized a stage when generally staggered Hox expression could be detected in ventral tissues (early trochophore 10 hpf, Fig. 4a). In both species, the staggered ventral Hox expression was along the AP axis according to the sequence of a presumptive Hox cluster (spatial collinearity) (Fig. 4b, c). We then found that the ventral tissues showing the most Hox expression in both species were neuroectodermal tissues because they expressed the pan-neural marker *soxb* (Fig. 4d). In *Ac. rubrolineata*, although a proportion of ventral signals were observed in subepidermal cells (Fig. 3n-w), these tissues are likely also neural tissues, because the tissues of its close relative *Ac. crinita* express the pan-neural marker *elav*^22^. Overall, staggered Hox expression was detected along the AP axis of the ventral neuroectoderm in both molluscan species, indicating that Hox genes play conserved roles in neurogenesis.

However, Hox expression in the neural tissues of *L. goshimai* lost its staggered pattern in as little as two hours (from 10 to 12 hpf, see Fig. 2 and supplemental figures S3-S4) and exhibited no obvious relationship with the AP axis in later larvae (supplemental figures S4-S5). This finding demonstrates that staggered Hox expression is restricted to the early phase of neurogenesis in *L. goshimai*. In *Ac. rubrolineata*, the staggered expression in ventral tissues was stable at the two stages examined (supplemental figures S6-S7).

### Dorsal Hox expression: tight correlation with the shell field

Although highly conserved staggered expression was detected in ventral tissues, dorsal Hox expression showed few common characteristics. Nevertheless, we noticed that dorsal Hox expression correlated with shell development in a lineage-specific manner. To provide comparable descriptions, we use the term “shell field” to refer to the whole area of the dorsal ectoderm that contributes to shell development, although this term may have had more specific meanings in previous literature^33^. In *L. goshimai*, the trochophore larva (10 hpf) possessed a single round shell plate (Fig. 5a). In accordance, *hox1-3* were expressed in a circular (or partially circular) pattern that surrounded the shell field (Fig. 5d-f). The other three genes, namely, *hox4*, *hox5* and *lox5*, were expressed in the central regions of the shell field (Fig. 5g-i). An association between the shell field and more Hox expression was detected in other developmental stages. In earlier stages (6- and/or 8-hpf gastrulae), the expression of *lox2* was detected in the dorsal ectoderm and thus may play a role in shell-field patterning (supplemental figures S1-S2). In the later trochophore larva (12 hpf), although the *hox3* expression in the shell field vanished, expression of *lox4*, *lox2* and *post2* in the leading edge of the shell field was evident (supplemental figure S4). In general, most Hox genes of *L. goshimai* showed a correlation with the shell field on the dorsal side at particular developmental stages.

In *Ac. rubrolineata*, in accordance with its larval shell field being composed of seven repeated shell plates/pseudosegments (Fig. 5b, c), *hox1-5* and *lox2* were expressed in particular bands in the dorsal ectoderm (Fig. 5j-o, j’-o’). For each of the other Hox genes, the expression covered a continuous region of the dorsal ectoderm, and a specific correlation between the expression and the shell field was difficult to determine (supplemental figures S6-S7). In summary, similar to the expression in *L. goshimai*, the dorsal Hox expression in *Ac. rubrolineata* also showed a tight correlation to the shell field, while the correlation was more specific for *hox1-5* and *lox2* and less specific for the other Hox genes.

## Discussion

### A generalized model of molluscan *Hox* expression

From our results, we deduce a generalized model of molluscan Hox expression based on dissociated dorsal and ventral expression. First, ventral Hox expression shows a conserved staggered pattern at particular developmental stages (i.e., during early neuroectoderm patterning). Second, dorsal Hox expression correlates with the shell field and is highly lineage-specific. Lastly, in later larval stages of conchiferans, most Hox genes are exclusively expressed in the nervous system (yet no longer in a staggered manner). By closely examining previous reports, we conclude that this generalized model is compatible with known Hox expression data from various molluscan clades. It is particularly evident in the scaphopod *Antalis entalis*, in which Hox expression at most developmental stages has been reported^23^. Hox expression in this species matches the generalized model, including the staggered ventral expression in early larvae (mid-trochophore), evident expression in the shell field at early stages, and exclusive neural expression in later larvae (in a non-staggered manner)^23^. In the polyplacophoran *Ac. crinita*, ventral expression in the neural tissues and dorsal expression in the shell field have also been described separately, although the dorsoventral dissociation was not emphasized^21,22^. Particular aspects of Hox expression have been reported in other species, which are actually snapshots of the generalized model, including the expression in the brachial crown (the modified foot) and the nervous system in the embryos of the cephalopod *Euprymna scolopes*^28^, the dorsal expression in the shell field of the gastrula in the bivalve *Patinopecten yessoensis*^24^, and the neural expression in the late larvae in the gastropod *Haliotis asinina*^25^. The only species whose Hox expression does not match the model is the gastropod *Gibbula varia*^26,27^. Given that Hox expression in *G. varia* shows very uncommon characteristics (*e.g.*, expression in the prototroch or velum^27^, which is never observed in other molluscs), further investigations on more developmental stages of *G. varia* and more gastropod species are needed to explain these unusual Hox expression patterns.

Based on these results, we propose that the relatively quick changes in molluscan Hox expression, in particular in conchiferans, have obscured the commonalities among different clades and that the dorsoventral dissociation further adds to the complexity. Although Hox expression has been described in much detail in previous studies, a generalized model has not been uncovered, likely because these studies focused on a single species and/or investigated limited developmental stages, making them insufficient for cross-species comparisons.

### Molluscan Hox genes: roles in neurogenesis and shell formation

The molluscan dorsal and ventral Hox expression patterns, despite their dissociation, indicate roles of Hox genes in neurogenesis and shell formation. The staggered expression in ventral neural tissues is reminiscent of the conserved staggered Hox expression in the nervous system of other bilaterian clades^9,34,35^. The potential roles of Hox genes in molluscan neurogenesis is consistent with a conserved mechanism of early neural patterning involving Hox genes in a wide range of animal lineages^36,37^. Moreover, because a large proportion of the neuroectoderm contributes to foot development, the roles of Hox genes in molluscan foot development are also indicated. On the dorsal side, although Hox expression in the shell field is frequently observed, the particular genes showing this expression pattern vary greatly among different clades^21,23,25,26^. While expression of most Hox genes can be detected in the shell field of the polyplacophoran *Ac. crinita*^21,22^, shell-field expression was limited to a small number of Hox genes in conchiferan molluscs (gastropod, bivalve and scaphopod)^23,24,26^. Here, we show that, similar to polyplacophorans, the expression of most Hox genes can be detected in the gastropod shell field at particular developmental stages. This result indicates that the involvement of Hox genes in shell-field patterning is far greater than previously realized and thus suggests a deep integration of Hox functions in molluscan shell development. The expression of more Hox genes in the shell field of other conchiferans is expected to be revealed as more developmental stages are investigated.

Lastly, although we use the general terms dorsal and ventral patterning for convenience, caution should be taken when discussing specific processes. Referring to these processes more generally as neural and non-neural patterning may be preferred. Posterior ectoderm patterning should also be paid special attention because, in particular species (e.g., *L. goshimai*), this region, unlike other ventral tissues, may not contain neuroectodermal tissues (indicated by the lack of *soxb* expression, see Fig. 4d) and does not exhibit the same obvious dorsoventral dissociation of Hox gene expression (Fig. 3k-l).

### The dissociated *Hox* expression in molluscs: evolutionary implications

Given that Hox genes play important roles in body patterning and that molluscan Hox expression shows both conserved and lineage-specific characteristics, this generalized Hox expression model provides useful information for inferring how diverse body plans evolved in molluscs. The expression data strongly suggest that the Hox genes contributed to patterning the nervous system (foot) ventrally and the shell field dorsally in the last common ancestor of molluscs. Nevertheless, regardless of the particular pattern (i.e., dorsal or ventral expression), we propose that the most important part is the dissociation itself because it allows the potential lineage-specific dorsal or ventral patterning, which may underlie the vast diversity of molluscan body plans. For instance, as we propose in Fig. 6, dorsal pseudosegments may have been formed in the lineage leading to polyplacophorans (assuming a non-segmented molluscan ancestor), and the dorsal part would have rotated in the lineage leading to gastropods. The unique shell shapes and structures in bivalves, scaphopods and cephalopods as well as the sclerites in aplacophorans represent other types of diversified dorsal structures (Fig. 6). Similarly, the diverse foot types in bivalves, scaphopods, cephalopods and aplacophorans might result from the lineage-specific patterning of the primitive ventral foot in the common molluscan ancestor that is likely still maintained in monoplacophorans, gastropods and polyplacophorans (Fig. 6). This lineage-specific ventral patterning does not contradict the conserved staggered Hox expression observed in early neurogenesis because this pattern can disappear quickly (e.g., in *L. goshimai*), and ventral Hox expression exhibits lineage-specific characteristics later in development. Together, the dissociated dorsal and ventral Hox expression, in combination with the robust involvement of Hox genes in morphogenesis, may underpin the diversification of the molluscan body plans.

**Fig. 6.**
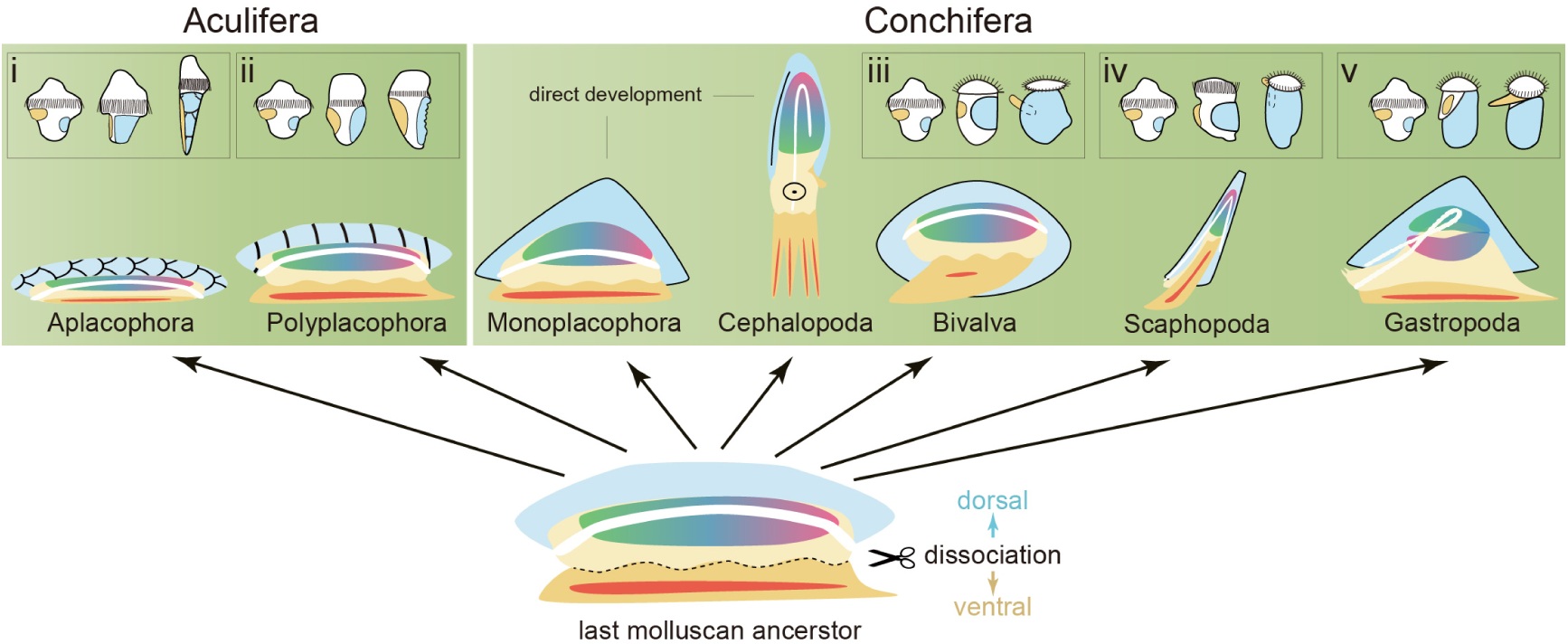
Evolutionary implications of dissociated dorsal and ventral patterning in molluscs. Spatially dissociated dorsal and ventral patterning may contribute to the diversity of the body plans characteristic of lineage-specific dorsal tissues (blue) and ventral tissues (orange). The nervous system (red) derived from the embryonic neuroectoderm is incorporated into the ventral tissues. Moreover, in conchiferans, the temporally dissociated dorsal and ventral patterning may allow earlier dorsal development and generate early larvae with large shells that can enclose the whole body (iii-v), which is not observed in aculiferans (i-ii). The diagrams of the larvae of aplacophorans and scaphopods are derived from previous studies^23,38^.

In addition to spatial dissociation, dorsal and ventral Hox expression also exhibited obvious temporal dissociation in *L. goshimai* (earlier dorsal expression). Similar earlier dorsal Hox expression has also been observed in a scaphopod^23^ and likely in a bivalve (solely dorsal expression at the early gastrula stage)^24^. Considering the speculated functions of Hox genes in the shell field and neuroectoderm/foot on the dorsal and ventral sides, respectively, the temporal dissociation of Hox expression suggests temporally dissociated dorsal and ventral patterning. Indeed, the earlier dorsal Hox expression in the three molluscan clades coincides with the bias towards earlier dorsal development in which the dorsal structure (shell field) develops earlier than the ventral structure (foot) (Fig. 6, inserts iii-v). In *L. goshimai*, this bias is reflected by *post1* expression that separates the dorsal and ventral tissues. When the shell field on the dorsal side expands quickly and encloses the larval body during the period from the gastrula to early veliger larvae, the ventral neuroectoderm/foot anlage remains small (supplemental figure S8). Such an earlier dorsal developmental strategy may provide survival advantages because it results in the quick formation of a large larval shell that can protect the larva from external threats. This developmental strategy seems to be common in indirectly developed conchiferans but not in aculiferans (compare inserts iii-v and i-ii in Fig. 6). The temporal dissociation of dorsal and ventral patterning represents a type of heterochrony and may contribute to the plasticity of developmental strategies in molluscan evolution.

### Common staggered hox expression in other spiralians

Similar to Hox expression in molluscs, Hox expression in other spiralians (except annelids) also shows diverse patterns and generally lacks staggered expression^10,11,16,17,39^. The demonstration of the dissociated Hox expression in dorsal and ventral tissues in molluscs provides a novel perspective for analysing spiralian Hox expression. Therefore, we re-analysed published data with the aim of recovering potential common characteristics (e.g., staggered expression) of spiralian Hox expression. When focusing on the ventral side, we recognized strikingly common staggered Hox expression at particular developmental stages of various spiralians, such as a rotifer^11^, nemertean^39^ and brachiopod^16^ (Fig. 7 and supplemental figures S9-S11). In annelids, although researchers have paid more attention to staggered Hox expression in segments, Hox expression is often observed in ventral tissues before segment formation^13,14,40,41^. To the best of our knowledge, the only two exceptions are the platyhelminth *Schmidtea*^10^ and the nemertean *Micrura*^17^, which may be due to the very unusual development of the animals (asexual reproduction in *Schmidtea* and the highly modified larval type in *Micrura*). As in molluscs, the ventral tissues of these spiralians showing staggered Hox expression are all neural tissues^11,13,14,16,39-41^ (in brachiopods, this idea was suggested subsequently by showing expression of neural patterning genes^42^). This result indicates that staggered Hox expression is largely maintained in various spiralians, which has long been debated. Similar to chordates^43^, combined Hox expression patterns including conserved staggered expression in AP patterning of neural tissues and extra expression in lineage-specific features seem to be widespread in spiralian lineages, indicating the roles of Hox genes in the evolution of the diverse spiralian body plans.

**Fig. 7.**
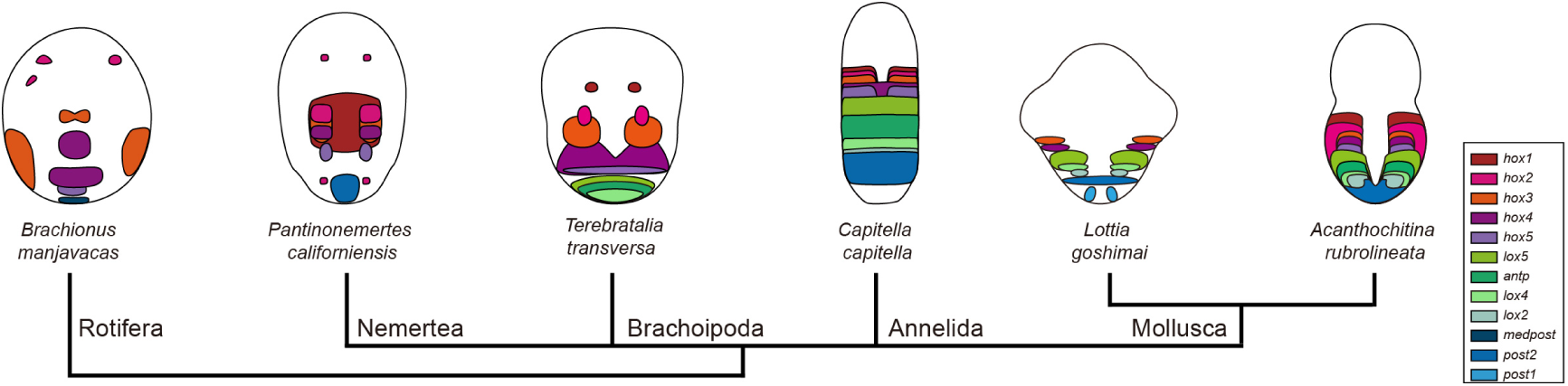
Staggered Hox expression in neural tissues in spiralians. The schematic diagrams of the rotifer, nemertean, brachiopod and annelid are derived from published data, and the details are provided in supplemental figures S9-S12.

## Methods

### Animals and larvae collection

Adults of *L. goshimai* Nakayama, Sasaki & Nakano, 2017 and *Ac. rubrolineata* (Lischke, 1873) were collected from intertidal rocks in Qingdao, China. Spawning occurred after the animals were transferred to the laboratory. For *L. goshimai*, which spawned relatively quickly after collection, each adult was placed into a single 100-ml cup for gamete collection. Artificial fertilization was conducted by mixing sperm and oocyte suspensions, and the fertilized eggs were cultured at 25°C. This procedure ensured the precise description of their developmental stages by referring to hpf. For *Ac. rubrolineata*, whose spawning was unpredictable, multiple individuals were put into the same container (20-25°C), and the embryos or larvae were collected on the second day. The developmental stages of the larvae were estimated based on their morphological characteristics. Trochophore larvae at two stages were used, which corresponded to the larvae ~24 hpf and ~48 hpf when cultured at 25°C (metamorphosis occurred 60-72 hpf). Specimens at the designated developmental stages of the two species (asterisks in Fig. 1b, c) were fixed in 4% paraformaldehyde in PBS containing 100 mM EDTA and 0.1% Tween-20 (pH 7.4). The veliger larvae of *L. goshimai* were anaesthetized by adding 1M magnesium chloride solution before fixation.

### Genes

Hox and SoxB genes were retrieved from the developmental transcriptomes of the two species by a BLAST search. Gene orthologies were verified by phylogenetic analysis (supplemental figures S13 and S15). For the Hox genes, characteristic residues of each orthologue were examined to confirm their orthologies as previously described^44,45^ (supplemental figure S14).

### In situ hybridization

Gene-specific primers containing a T7 promoter sequence (taatacgactcactataggg) were used to amplify the cDNA fragment of each gene. The resultant PCR products were used as templates in subsequent in vitro transcription to synthesize the digoxigenin-labelled probes. In situ hybridization was performed as previously described^24^ except that, after specimen rehydration, an additional protease K treatment was performed (20 min at room temperature with continuous shaking) before the incubation in 1% triethanolamine. Different concentrations of protease K (diluted in PBST) were used for different specimens: 25 μg/ml for *L. goshimai* embryos (8 hpf and before) and early larva (10 hpf), 50 μg/ml for *L. goshimai* mid-larvae (12 and 14 hpf) and all *Ac. rubrolineata* larvae, 75 μg/ml for *L. goshimai* late larvae (17 and 28 hpf).

### Scanning electronic microscopy

The specimens were fixed in 2.5% glutaraldehyde at 4°C overnight and then dehydrated in 100% ethanol. Drying, coating and observations under a scanning electronic microscope were performed as described previously^46^.

## Data availability

*L. goshimai* and *Ac. rubrolineata* sequence data generated during the current study have been deposited in GenBank with the primary accession numbers <u>MK637053</u> to <u>MK637075</u>.

## Acknowledgements

We thank Peter W. H. Holland for thoughtful discussions and critical revisions of the manuscript. Sincere thanks are due to Andreas Hejnol, Gregory A. Wray, Fei Xu and Lingyu Wang for suggestions on the manuscript. We thank Junlong Zhang for assistance in sample collection and species identification and for providing some of the included images of molluscs. We thank Weihong Yang for performing some of the in situ hybridization experiments. The study was funded by National Key R&D Program of China (2018YFD0900104) grant to P.H., the Marine S&T Fund of Shandong Province for Pilot National Laboratory for Marine Science and Technology (Qingdao) (2018SDKJ0302-1) grant to B.L., the China Agriculture Research System (CARS-49) grant to B.L. and the National Natural Science Foundation of China (41776157) grant to P.H.

## Author contributions

B.L. and P.H. conceived the project. P.H. collected animals, cloned Hox genes and interpreted the primary data, including concluding the generalized molluscan Hox expression model, proposing the hypothesis of molluscan body-plan diversification and perceiving the staggered Hox expression in non-mollusc spiralians. Q.W. performed the in situ hybridization experiments, record raw images, cloned SoxB genes and contributed to animal collection. P.H. and Q.W. prepared the figures. S.T. contributed to the experiment design by speculating dynamic Hox expression. P.H. wrote the manuscript. B.L. oversaw the whole project, contributed to data analysis and critically revised the manuscript. All authors discussed the results and commented on the manuscript.

## References

1 Garcia-Fernàndez, J. The genesis and evolution of homeobox gene clusters. Nature Reviews Genetics 6, 881–892, doi:10.1038/nrg1723 (2005).

2 Pearson, J. C., Lemons, D. & McGinnis, W. Modulating Hox gene functions during animal body patterning. Nature Reviews Genetics 6, 893–904, doi:10.1038/nrg1726 (2005).

3 Holland, P. W. H. Evolution of homeobox genes. Wiley Interdisciplinary Reviews: Developmental Biology 2, 31–45, doi:10.1002/wdev.78 (2013).

4 Dubuc, T. Q., Stephenson, T. B., Rock, A. Q. & Martindale, M. Q. Hox and Wnt pattern the primary body axis of an anthozoan cnidarian before gastrulation. Nature Communications 9, doi:10.1038/s41467-018-04184-x (2018).

5 Kroesen, A. E. et al. An axial Hox code controls tissue segmentation and body patterning in *Nematostella vectensis*. Science 361, 1377–1380, doi:10.1126/science.aar8384 (2018).

6 Cj, K. et al. The dance of the Hox genes - Patterning the anteroposterior body axis of *Caenorhabditis elegans*. Cold Spring Harbor Symposia on Quantitative Biology 62, 293–305 (1998).

7 Aronowicz, J. & Lowe, C. J. Hox gene expression in the hemichordate *Saccoglossus kowalevskii* and the evolution of deuterostome nervous systems. Integrative and Comparative Biology 46, 890–901, doi:10.1093/icb/icl045 (2006).

8 Tsuchimoto, J. & Yamaguchi, M. Hox expression in the direct-type developing sand dollar *Peronella japonica*. Developmental Dynamics 243, 1020–1029, doi:10.1002/dvdy.24135 (2014).

9 Hejnol, A. & Martindale, M. Q. Coordinated spatial and temporal expression of Hox genes during embryogenesis in the acoel *Convolutriloba longifissura*. BMC Biology 7, 65–65, doi:10.1186/1741-7007-7-65 (2009).

10 Currie, K. W. et al. HOX gene complement and expression in the planarian *Schmidtea mediterranea*. EvoDevo 7, 7–7, doi:10.1186/s13227-016-0044-8 (2016).

11 Fröbius, A. C. & Funch, P. Rotiferan Hox genes give new insights into the evolution of metazoan bodyplans. Nature Communications 8, doi:10.1038/s41467-017-00020-w (2017).

12 Papillon, D., Perez, Y., Fasano, L., Le Parco, Y. & Caubit, X. Hox gene survey in the chaetognath Spadella cephaloptera: evolutionary implications. Development Genes and Evolution 213, 142–148, doi:10.1007/s00427-003-0306-z (2003).

13 Kulakova, M. et al. Hox gene expression in larval development of the polychaetes Nereis virens and Platynereis dumerilii (Annelida, Lophotrochozoa). Development Genes and Evolution 217, 39–54, doi:10.1007/s00427-006-0119-y (2006).

14 Fröbius, A. C., Matus, D. Q. & Seaver, E. C. Genomic organization and expression demonstrate spatial and temporal Hox gene colinearity in the lophotrochozoan *Capitella sp. I*. PLoS ONE 3, doi:10.1371/journal.pone.0004004 (2008).

15 Irvine, S. Q. & Martindale M. Q. Expression patterns of anterior Hox genes in the polychaete *chaetopterus*: Correlation with morphological boundaries. Developmental Biology 217, 333–351, doi:10.1006/dbio.1999.9541 (2000).

16 Schiemann, S. M. et al. Clustered brachiopod Hox genes are not expressed collinearly and are associated with lophotrochozoan novelties. Proceedings of the National Academy of Sciences 114, E1913–E1922, doi:10.1073/pnas.1614501114 (2017).

17 Hiebert, L. S. & Maslakova, S. A. Hox genes pattern the anterior-posterior axis of the juvenile but not the larva in a maximally indirect developing invertebrate, *Micrura alaskensis* (Nemertea). BMC Biology 13, doi:10.1186/s12915-015-0133-5 (2015).

18 Smith, S. A. et al. Resolving the evolutionary relationships of molluscs with phylogenomic tools. Nature 480, 364–367, doi:10.1038/nature10526 (2011).

19 Haszprunar, G. & Wanninger, A. Molluscs. Current Biology 22, R510–R514, doi:10.1016/j.cub.2012.05.039 (2012).

20 Wanninger, A. & Wollesen, T. The evolution of molluscs. Biological Reviews 94, 102–115, doi:10.1111/brv.12439 (2019).

21 Fritsch, M., Wollesen, T., de Oliveira, A. L. & Wanninger, A. Unexpected co-linearity of Hox gene expression in an aculiferan mollusk. BMC Evolutionary Biology 15, doi:10.1186/s12862-015-0414-1 (2015).

22 Fritsch, M., Wollesen, T. & Wanninger, A. Hox and ParaHox gene expression in early body plan patterning of polyplacophoran mollusks. Journal of Experimental Zoology Part B: Molecular and Developmental Evolution 326, 89–104, doi:10.1002/jez.b.22671 (2016).

23 Wollesen, T., Rodríguez Monje, S. V., de Oliveira, A. L. & Wanninger, A. Staggered Hox expression is more widespread among molluscs than previously appreciated. Proceedings of the Royal Society B 285, 20181513–20181513, doi:10.1098/rspb.2018.1513 (2018).

24 Wang, S. et al. Scallop genome provides insights into evolution of bilaterian karyotype and development. Nature Ecology & Evolution 1, 0120–0120, doi:10.1038/s41559-017-0120 (2017).

25 Hinman, V. F., O’Brien, E. K., Richards, G. S. & Degnan, B. M. Expression of anterior Hox genes during larval development of the gastropod *Haliotis asinina*. Evolution and Development 5, 508–521, doi:10.1046/j.1525-142X.2003.03056.x (2003).

26 Samadi, L. & Steiner, G. Involvement of Hox genes in shell morphogenesis in the encapsulated development of a top shell gastropod (*Gibbula varia* L.). Development Genes and Evolution 219, 523–530, doi:10.1007/s00427-009-0308-6 (2009).

27 Samadi, L. & Steiner, G. Expression of Hox genes during the larval development of the snail, *Gibbula varia* (L.)-further evidence of non-colinearity in molluscs. Development Genes and Evolution 220, 161–172, doi:10.1007/s00427-010-0338-0 (2010).

28 Lee, P. N., Callaerts, P., De Couet, H. G. & Martindale, M. Q. Cephalopod Hox genes and the origin of morphological novelties. Nature 424, 1061–1065, doi:10.1038/nature01872 (2003).

29 Henry, J. J. & Martindale, M. Q. Conservation and innovation in spiralian development. Hydrobiologia 402, 255–265, doi:10.1023/a:1003756912738 (1999).

30 Hejnol, A. A twist in time-the evolution of spiral cleavage in the light of animal phylogeny. Integrative and Comparative Biology 50, 695–706, doi:10.1093/icb/icq103 (2010).

31 Lambert, J. D. Developmental Patterns in Spiralian Embryos. Current Biology 20, R72–R77, doi:10.1016/j.cub.2009.11.041 (2010).

32 Nielsen, C. Some aspects of spiralian development. Acta Zoologica 91, 20–28, doi:10.1111/j.1463-6395.2009.00421.x (2010).

33 Jackson, D. J. & Degnan, B. M. The importance of evo-devo to an integrated understanding of molluscan biomineralisation. Journal of Structural Biology 196, 67–74, doi:10.1016/j.jsb.2016.01.005 (2016).

34 Kourakis, M. J. et al. Conserved anterior boundaries of Hox gene expression in the central nervous system of the leech *Helobdella*. Developmental Biology 190, 284–200, doi:10.1006/dbio.1997.8689 (1997).

35 Serano, J. M. et al. Comprehensive analysis of Hox gene expression in the amphipod crustacean *Parhyale hawaiensis*. Developmental Biology 409, 297–309, doi:10.1016/j.ydbio.2015.10.029 (2016).

36 Arendt, D., Tosches, M. A. & Marlow, H. From nerve net to nerve ring, nerve cord and brain-evolution of the nervous system. Nature Reviews Neuroscience 17, 61–72, doi:10.1038/nrn.2015.15 (2016).

37 Hartenstein, V. & Stollewerk, A. The evolution of early neurogenesis. Developmental Cell 32, 390–407, doi:10.1016/j.devcel.2015.02.004 (2015).

38 Todt, C. & Wanninger, A. Of tests, trochs, shells, and spicules: Development of the basal mollusk *Wirenia argentea* (Solenogastres) and its bearing on the evolution of trochozoan larval key features. Frontiers in zoology 7, 6–6, doi:10.1186/1742-9994-7-6 (2010).

39 Hiebert, L. S. & Maslakova, S. A. Expression of Hox, Cdx, and Six3/6 genes in the hoplonemertean *Pantinonemertes californiensis* offers insight into the evolution of maximally indirect development in the phylum Nemertea. EvoDevo 6, 1–15, doi:10.1186/s13227-015-0021-7 (2015).

40 Irvine, S. Q. & Martindale, M. Q. Comparative analysis of hox gene expression in the polychaete *Chaetopterus*. Library 651, 640–651 (2001).

41 Kourakis, M. J. & Martindale, M. Q. Hox gene duplication and deployment in the annelid leech *Helobdella*. Evolution and Development 3, 145–153, doi:10.1046/j.1525-142X.2001.003003145.x (2001).

42 Martín-Durán, J. M. et al. Convergent evolution of bilaterian nerve cords. Nature 553, 45–50, doi:10.1038/nature25030 (2018).

43 Lappin, T. R. J., Grier, D. G., Thompson, A. & Halliday, H. L. HOX genes: Seductive science, mysterious mechanisms. Ulster Medical Journal 75, 23–31 (2006).

44 Balavoine, G., De Rosa, R. & Adoutte, A. Hox clusters and bilaterian phylogeny. Molecular Phylogenetics and Evolution 24, 366–373, doi:10.1016/s1055-7903(02)00237-3 (2002).

45 Pérez-Parallé, M. L., Pazos, A. J., Mesías-Gansbiller, C. & Sánchez, J. L. Hox, Parahox, Ehgbox, and NK Genes in Bivalve Molluscs: Evolutionary Implications. Journal of Shellfish Research 35, 179–190, doi:10.2983/035.035.0119 (2016).

46 Tan, S., Huan, P. & Liu, B. Expression patterns indicate that BMP2/4 and Chordin, not BMP5-8 and Gremlin, mediate dorsal-ventral patterning in the mollusk *Crassostrea gigas*. Development Genes And Evolution 227, 75–84 (2017).

