## Supplementary materials for "Dorsoventral dissociation of Hox gene expression underpins the diversification of molluscs"

**Supplemental figures S1-S15**

**Supplemental Table S1-S2**

**Supplemental references (1-7)**

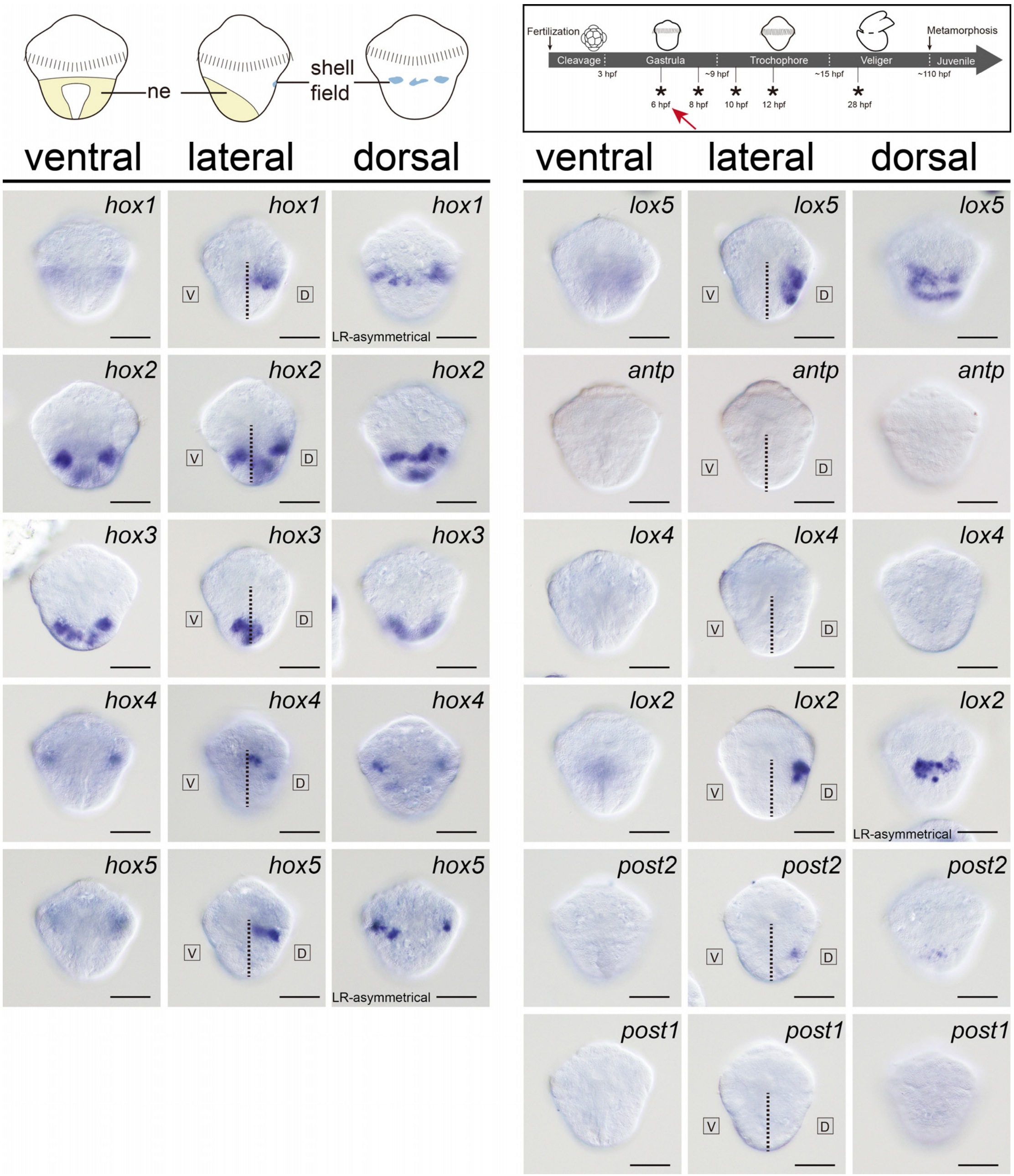


**Supplemental figure S1 Hox expression in the 6-hpf gastrula of *L. goshimai*.** At this stage, gastrulation is ongoing and the dorsal-ventral axis is actually not fully established. The blastopore is not formed either, while the shell field is only on its initial formation. Despite the still ongoing dorsal-ventral patterning, we still use the dashed lines to separate the "dorsal" (D) and "ventral" (V) tissues for a purpose of comparison. Expression of most Hox genes were detected except *antp*, *lox4* and *post1*. Almost all Hox expression is exclusively dorsal, while *hox2* show additional ventral expression. Another exception is *hox3*, whose expression was exclusively ventral. However, such "ventral" expression of *hox3* actually emerged at dorsal side firstly, which moved to ventral side during gastrulation (data not shown). Some genes show left-right asymmetrical (LR-asymmetrical) expression. ne: neuroectoderm. Bars represent 50 µm.

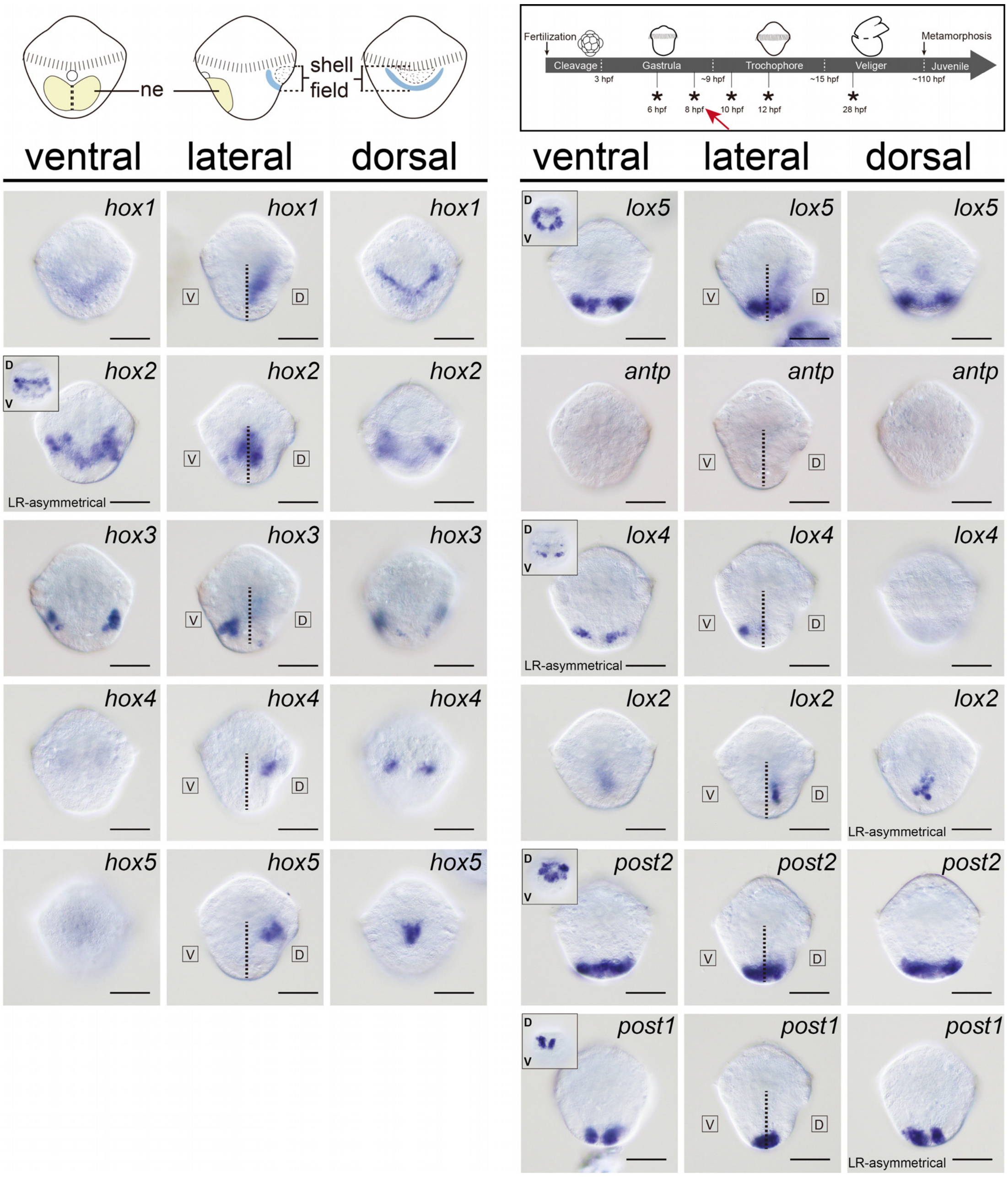


**Supplemental figure S2 Hox expression in the 8-hpf gastrula of *L. goshimai*.** The dashed lines separate the dorsal (D) and ventral (V) tissues. The small inserts in some panels show the posterior views. At this stage, gastrulation is about to finish. On the dorsal side, the central region of the shell field exhibits numerous lamellipodia, indicating the shell plate will form soon. The neuroectoderm can be well discriminated on the ventral side. Expression of all Hox genes except *antp* were detected. Although most Hox expression was still dorsal, however, the expression patterns change significantly comparing to that at the previous stage (6 hpf). Two genes (*lox5* and *post2*) show circumferential expression. The only gene showing exclusively ventral expression is *lox4*. The expression of *post1* is detected in the terminal of the embryo. Some genes show left-right asymmetrical (LR-asymmetrical) expression. ne: neuroectoderm. Bars represent 50 μm.

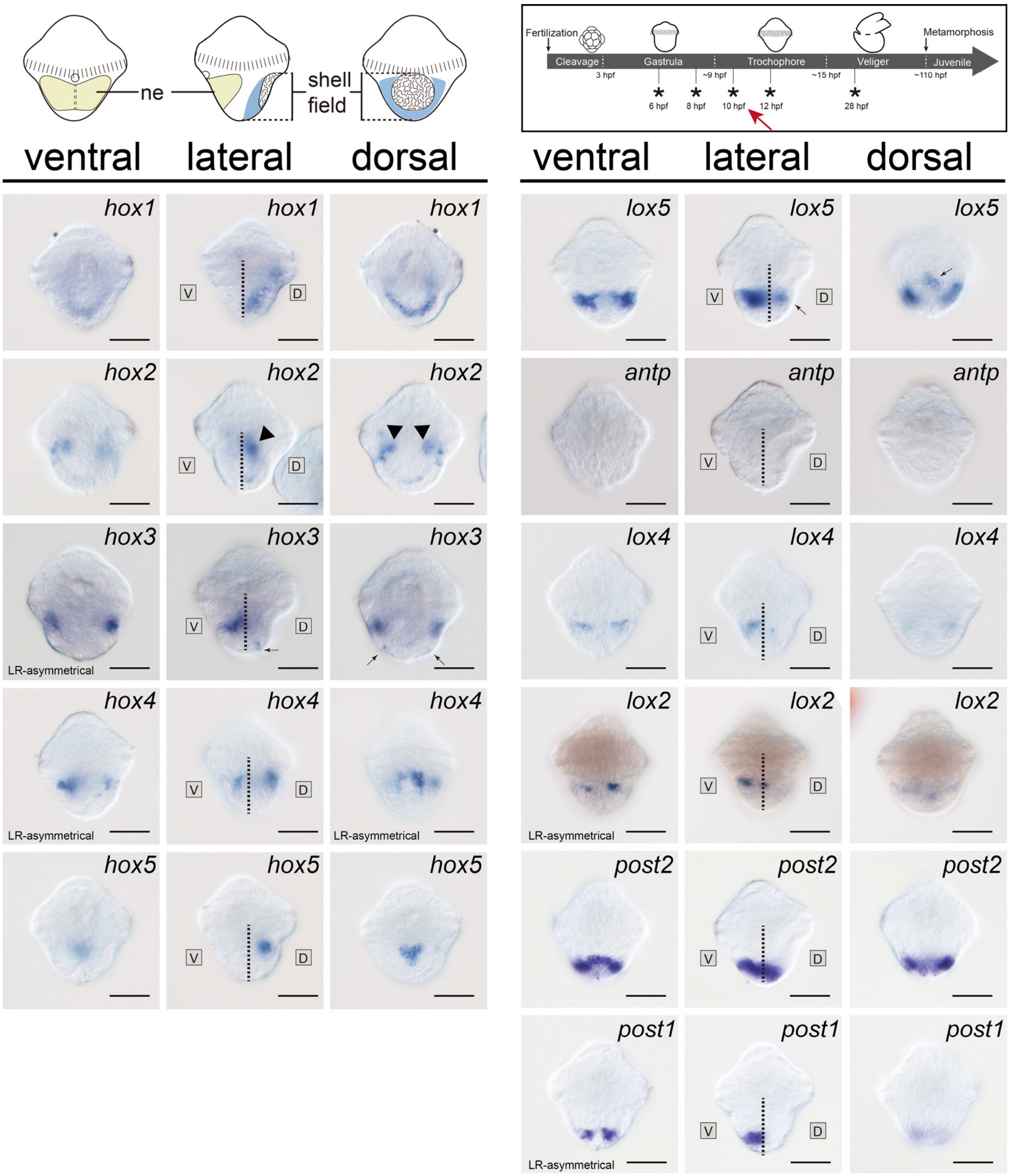


**Supplemental figure S3 Hox expression in the 10-hpf trochophore larva of *L. goshimai*.** The dashed lines separate the dorsal (D) and ventral (V) tissues. At this stage, a single round shell plate can be observed on the dorsal side, and the bilaterally symmetrical neuroectoderm can be easily discriminated on the ventral side. Although the number of Hox genes showing dorsal expression does not change significantly, the expression levels decrease generally. On the contrary, ventral expression was detected for much more genes comparing to 8-hpf gastrula. The majority of *hox2* expression was detected in subepidermal cells, which was likely mesodermal expression (black arrowheads). Moreover, the ventral Hox expression showed a generally staggered pattern (see Fig. 4). The ventral tissues expressing Hox genes also express the pan-neural marker *soxb*, indicating they were the neuroectodermal tissues (see Fig. 4). Note that because neuroectodermal tissues spread to the lateral sides, the expression of some anterior Hox genes (*hox3* and *hox4*) was actually observed in lateral tissues. For convenience, however, we still described it to be ventral expression. On the dorsal side, *hox1-5* all showed correlation with the round shell plate. Among them, *hox1-3* showed circular (or partially circular) expression surrounding the shell plate, and *hox4-5* showed expression in the central region of the shell plate. Another gene, *lox5*, also showed central expression in the shell plate similar to that of *hox5*. Some genes show left-right asymmetrical (LR-asymmetrical) expression. ne: neuroectoderm. Bars represent 50 μm.

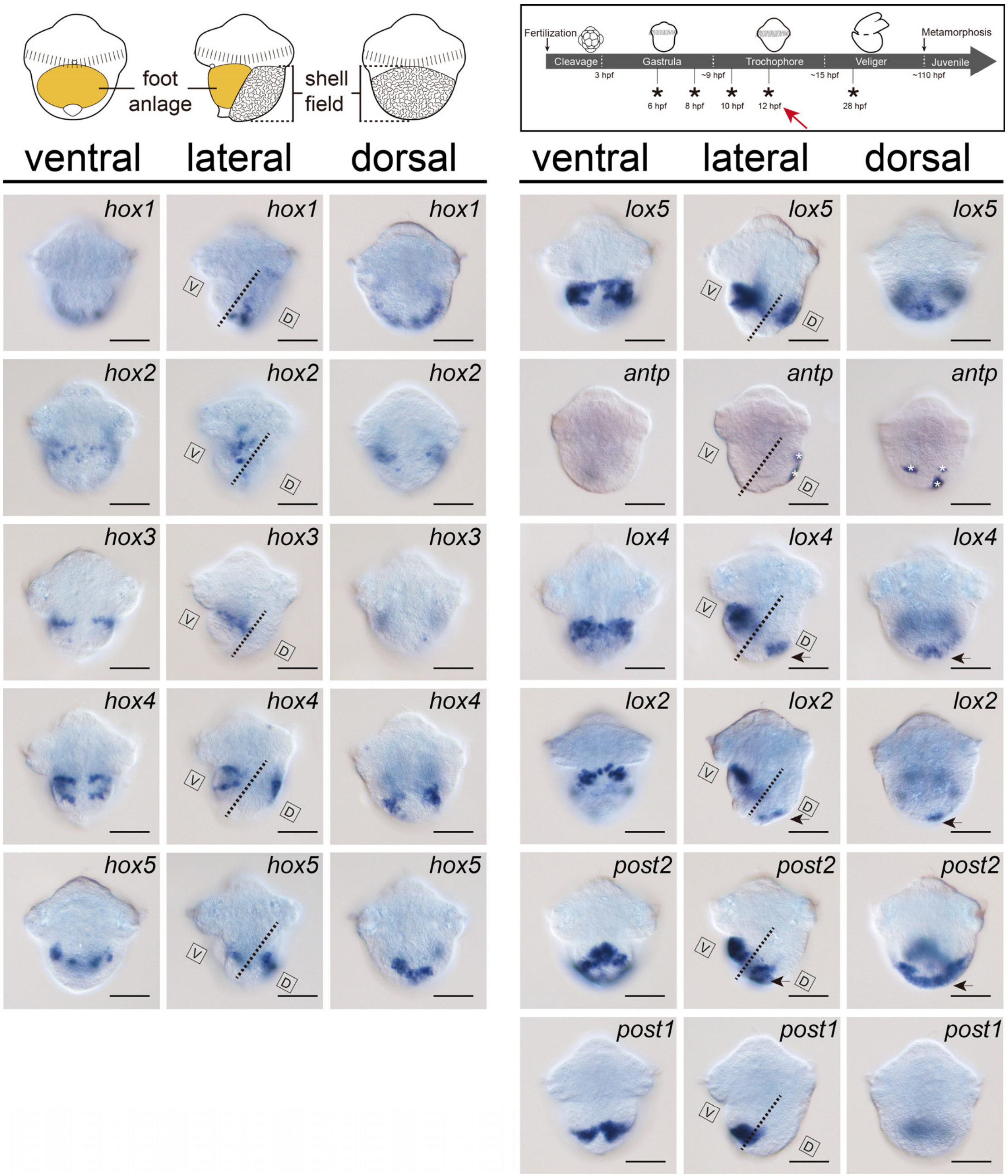


**Supplemental figure S4 Hox expression in the 12-hpf trochophore larva of *L. goshimai*.** The dashed lines separate the dorsal (D) and ventral (V) tissues, which become tilted due to the expansion of shell field. Compared to the 10-hpf larva, *hox3* expression in the shell field disappeared, whereas expression of *lox4*, *lox2* and *post2* emerged in the leading edge of the shell field (black arrows). On the ventral side, the foot anlage is formed, in which the majority of the neuroectoderm is involved. All Hox genes except *hox1* and *antp* show expression in the foot anlage at relatively high levels. However, the ventral expression no longer exhibits a staggered pattern. White asterisks indicate unspecific staining. Bars represent 50 µm

.

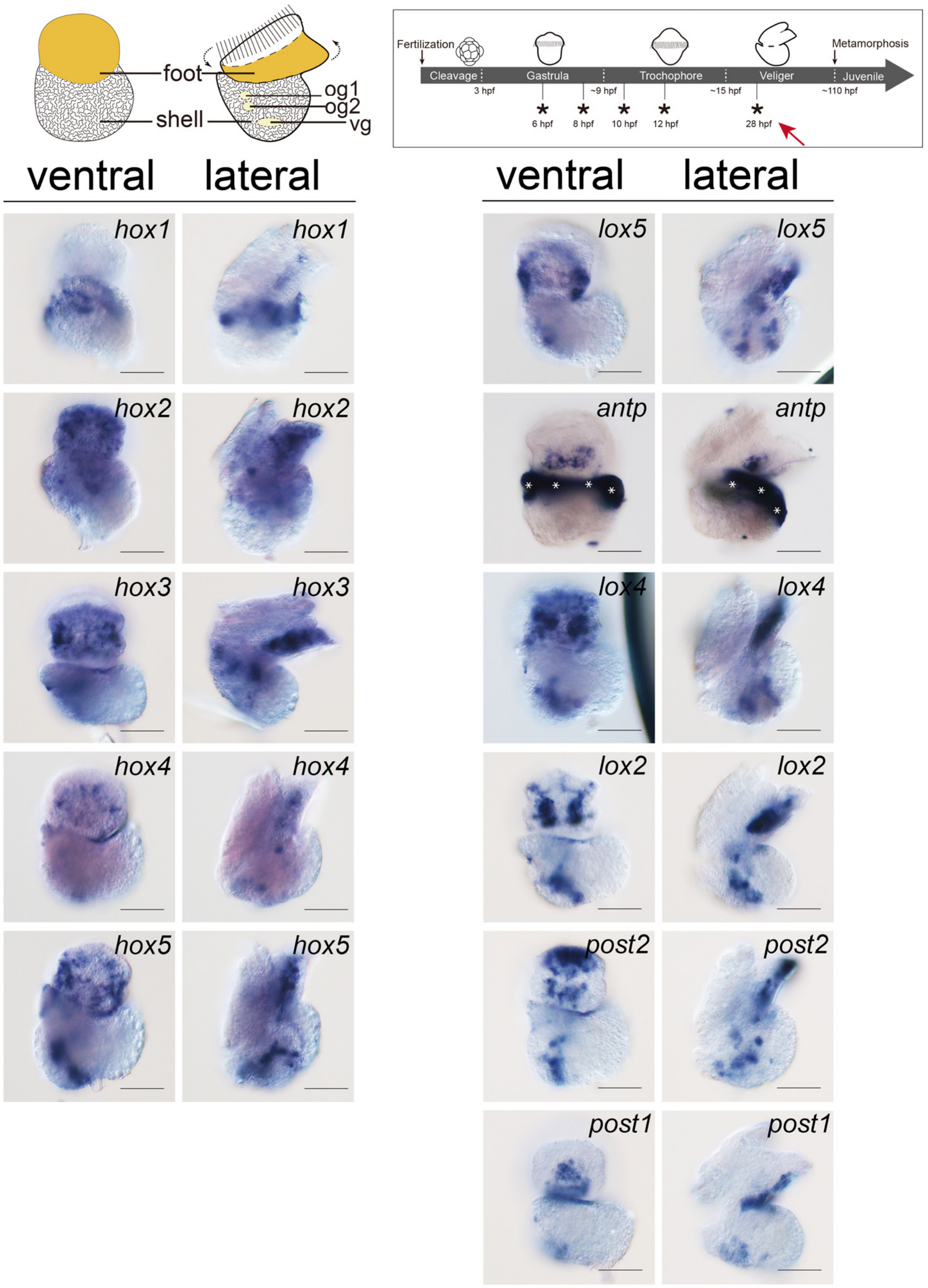


**Supplemental figure S5 Hox expression in the 28-hpf veliger larva of *L. goshimai*.** At this stage, the well-developed larval shell is able to enclose the larval body and it is not possible to separate dorsal and ventral tissues simply by a straight line. A larval foot is formed. However, it now faces to the opposite side (i.e., the dorsal side at the previous stages), which is caused by torsion (dashed arrows). Hox expression was detected in exclusively neural tissues of the foot (pedal ganglia) and internal organs [supraesophageal (og1), subesophageal (og2) and visceral ganglions(vg)]. As an only exception, *hox1* still showed major expression in the mantle tissues surrounding the shell edge, although weak expression was also observed in the foot. White asterisks indicate unspecific staining. Bars represent 50 µm.

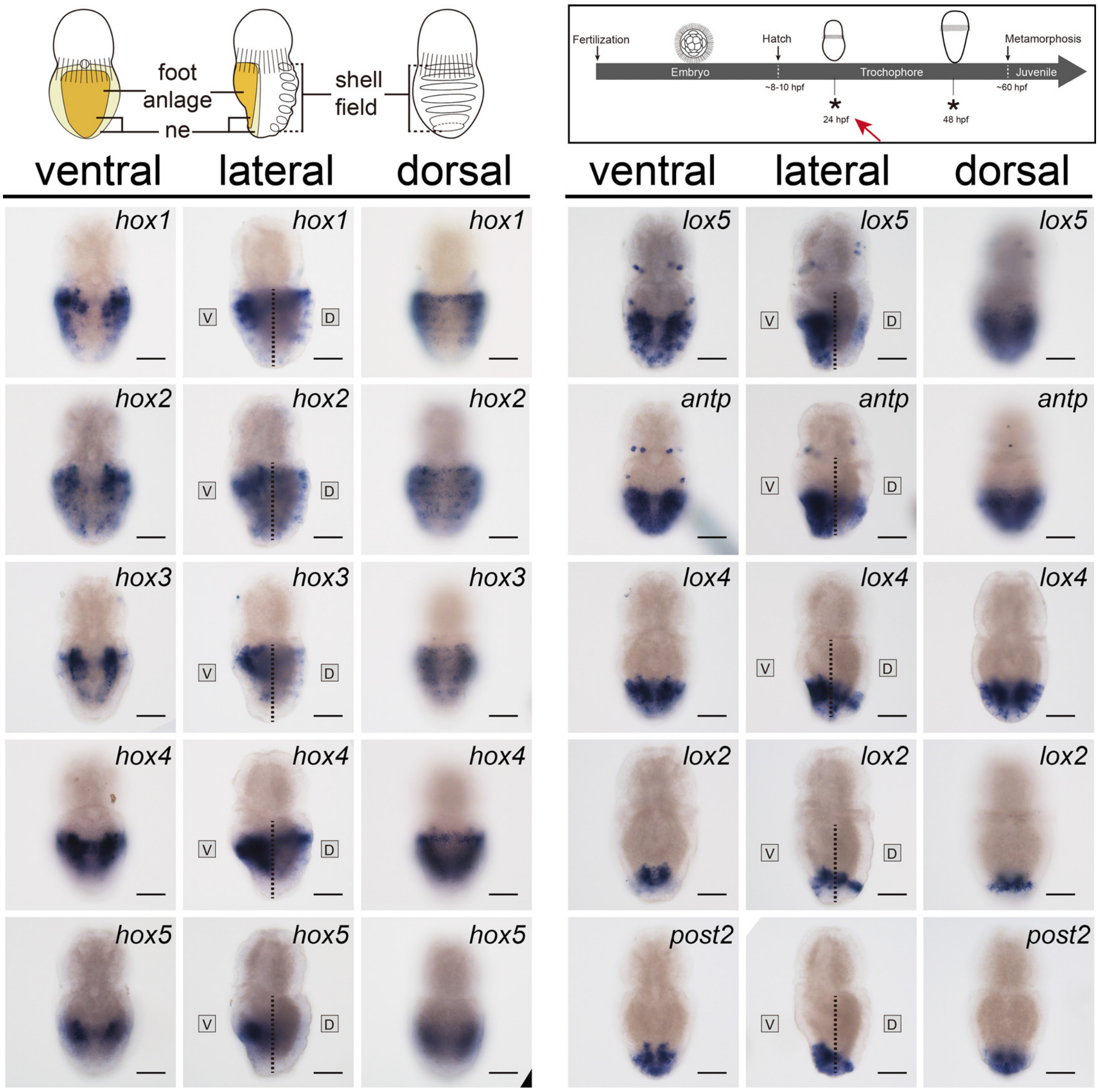


**Supplemental figure S6 Hox expression in the 24-hpf trochophore larva of *Ac. rubrolineata.*** The dashed lines separate the dorsal (D) and ventral (V) tissues. Six to seven pseudosegments can be discriminated on the dorsal side (see Fig. 5b) and a foot anlage covered by cilia is developed on the ventral side (not shown). All Hox expression is detected in both dorsal and ventral tissues except *hox5* showing exclusive ventral expression. The dorsal and ventral expression is markedly different for most genes. ne: neuroectoderm. Bars represent 50 µm.

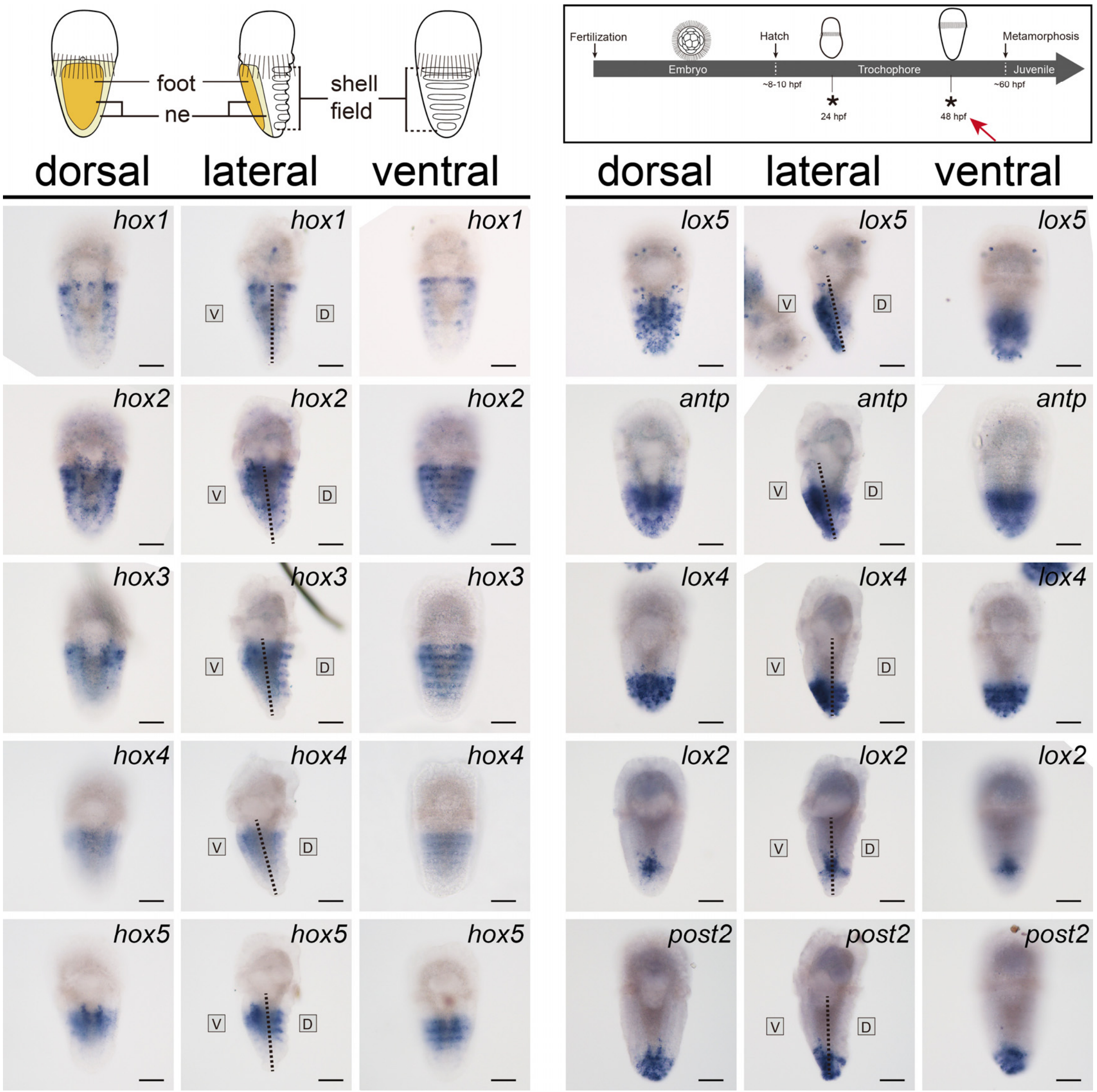


**Supplemental figure S7 Hox expression in the 48-hpf trochophore larva of *Ac. rubrolineata*.** The dashed lines separate the dorsal (D) and ventral (V) tissues, some of which tilted because the larvae bend to the dorsal side (may be caused by fixation). Multiple secretory papillae can be observed on the surface of the seven pseudosegments on the dorsal side (see Fig. 5c), indicating shell plates will form soon at the sites. A foot can be observed on the ventral side (not shown). The Hox expression does not change significantly comparing to the 24-hpf larva. With the development of pseudosegments, the dorsal Hox expression in the pseudosegment becomes more obvious. Among them, expression of *hox1-5* and *lox2* showed obvious correlation with particular pseudosegments, while the dorsal expression of other genes covers particular regions and the correlation with the shell field is not so specific. The staggered pattern on the ventral side sustained at this stage. ne: neuroectoderm. Bars represent 50 μm.

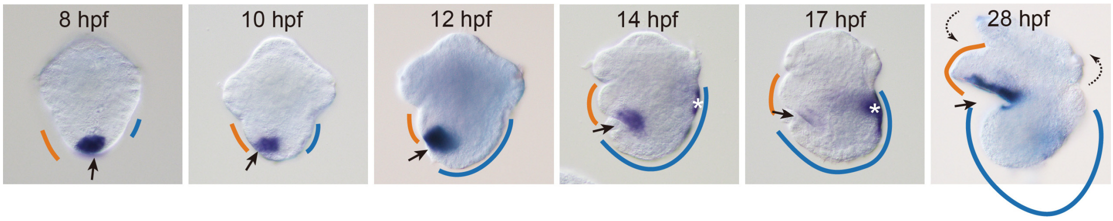


**Supplemental figure S8 The continuous migration of *post1*-expressing tissue during early development of *L. goshimai*.** The terminal tissue expressing *post1* (black arrows) migrate ventrally during the development from gastrula (8 hpf) to veliger larva (28 hpf). During this period, the dorsal shell field (blue lines) expanded quickly while the ventral tissues (mainly refer to the neuroectoderm, which later develops into larval foot, indicated by the orange lines) remain small. Note that, due to torsion (dashed arrows), the anterior part of the 28-hpf larva (mainly refers to the velum and foot) actually faces the opposite side comparing to other embryos/larvae. However, here we still put the larva with foot toward left like other larvae for the convenience of comparison. White asterisks indicate non-specific staining.

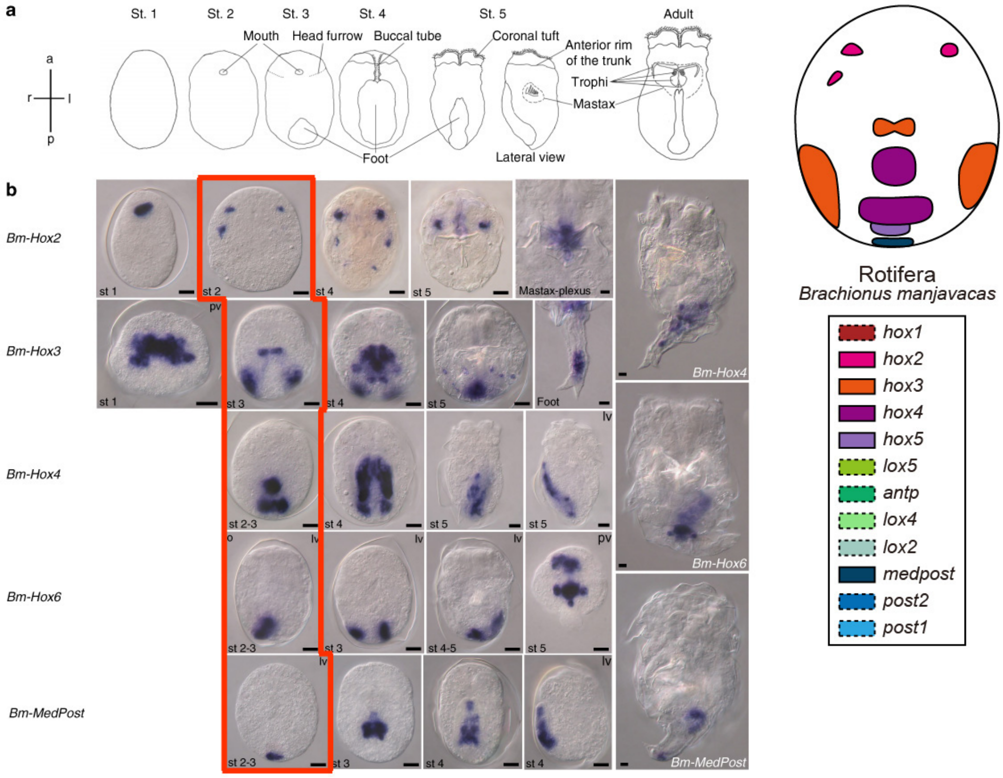


**Fig. 2** Expression of *Hox* genes during embryogenesis of *Brachionus manjavacas*. **a** Schematic of embryonic stages of *Brachionus manjavacas* with morphological characteristics used for staging. **b** Whole-mount in situ hybridization on amictic female embryos. Adults are only shown for genes with expression persisting into the adult stage. Anterior to the top. Mostly ventral views are shown. pv, posterior view, dorsal side up; lv, lateral view, ventral to the left. Scale bar, 10 µm

**Supplemental figure S9 The generally staggered Hox expression in the ventral tissues of the stage2-3 embryo of the rotifer *Brachionus manjavacas*.** The original figure is published previously (Figure 2 in [1]) and reproduced here. A schematic diagram derived from the ISH results enclosed by the red frame is shown and also included in Fig. 7.

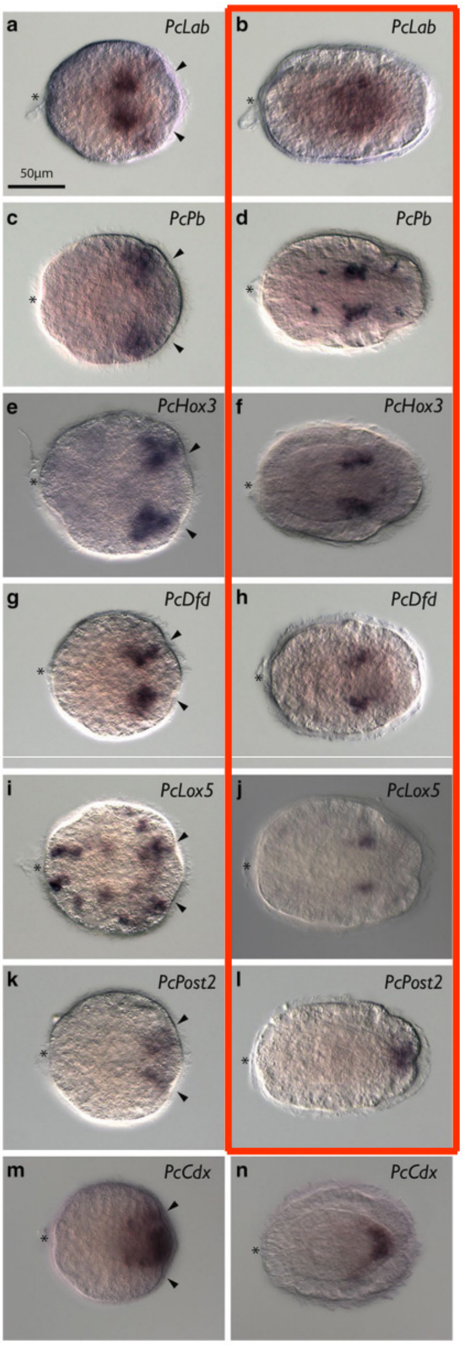


**Fig. 3** *Hox* and *Cdx* expression in *P. californiensis*. All images show frontal views with larval anterior to the *left*. *Left column* shows invagination-stage larvae. *Right column* shows rudiment-stage larvae. *Arrowheads* indicate paired posterior invaginations. *Asterisk* indicates apical organ.

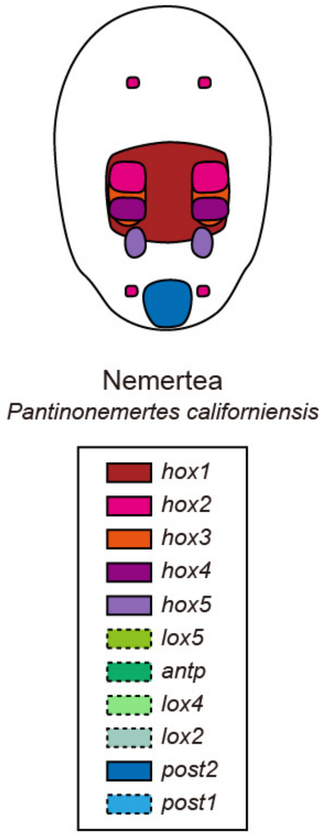


**Supplemental figure S10 The generally staggere**d Hox exp**ression in the ventral tissues of the pilidium larva of the nemertean *Micrura alaskensis*.** The original figure is published previously (Figure 3 in [2]) and reproduced here. A schematic diagram derived from the ISH results enclosed by the red frame is shown and also included in Fig. 7.

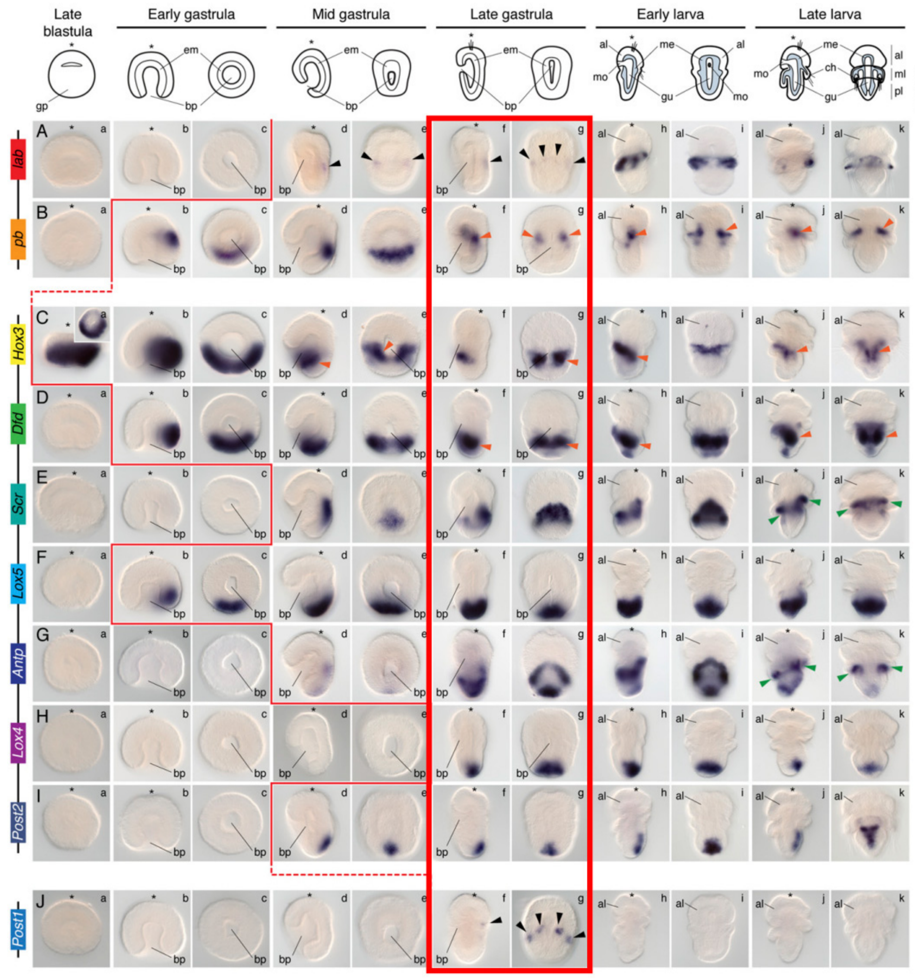


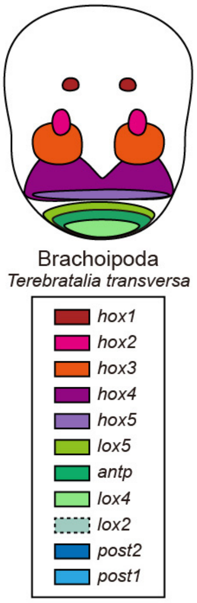


**Fig. 2.** Whole-mount in situ hybridization of each Hox gene during embryonic and larval stages in *T. transversa*. The Hox genes *lab* and *Post1* are expressed during chaetae formation. The genes *pb*, *Hox3*, and *Dfd* are expressed collinearly along the mantle and pedicle mesoderm. The Hox genes *Scr* and *Antp* are expressed in the periostracum, the shell-forming epithelium. *Lox5*, *Lox4*, and *Post2* are expressed in the posterior ectoderm of the pedicle lobe. These expression patterns are described in detail in the text. Black arrowheads indicate expression in the chaetae sacs. Orange arrowheads highlight mesodermal expression Green arrowheads indicate expression in the periostracum. The genomic organization of the Hox genes is shown on the left. On top are schematic representations of each analyzed developmental stage on its respective perspective. In these schemes, the blue area represents the mesoderm. Drawings are not to scale. The red line indicates the onset of expression of each Hox gene based on in situ hybridization data. The blastula stage is a lateral view (*Inset* in Ca is a vegetal view). For each other stage, the left column is a lateral view and the right column is a dorsoventral view. The asterisk demarcates the animal/anterior pole. al, apical lobe; bp, blastopore; ch, chaetae; em, endomesoderm; gp, gastral plate; gu, gut; me, mesoderm; ml, mantle lobe; mo, mouth; pl, pedicle lobe

**Supplemental figure S11 The generally staggered Hox expression in the ventral tissues of the late gastrula of the brachiopod *Terebratalia transversa*.** The original figure is published previously (Figure 2 in [3]) and reproduced here with permission. A schematic diagram derived from the ISH results enclosed by the red frame is shown and also included in Fig. 7. Note that the colors we used to represent the Hox genes in the diagram are different from those used in the original article.

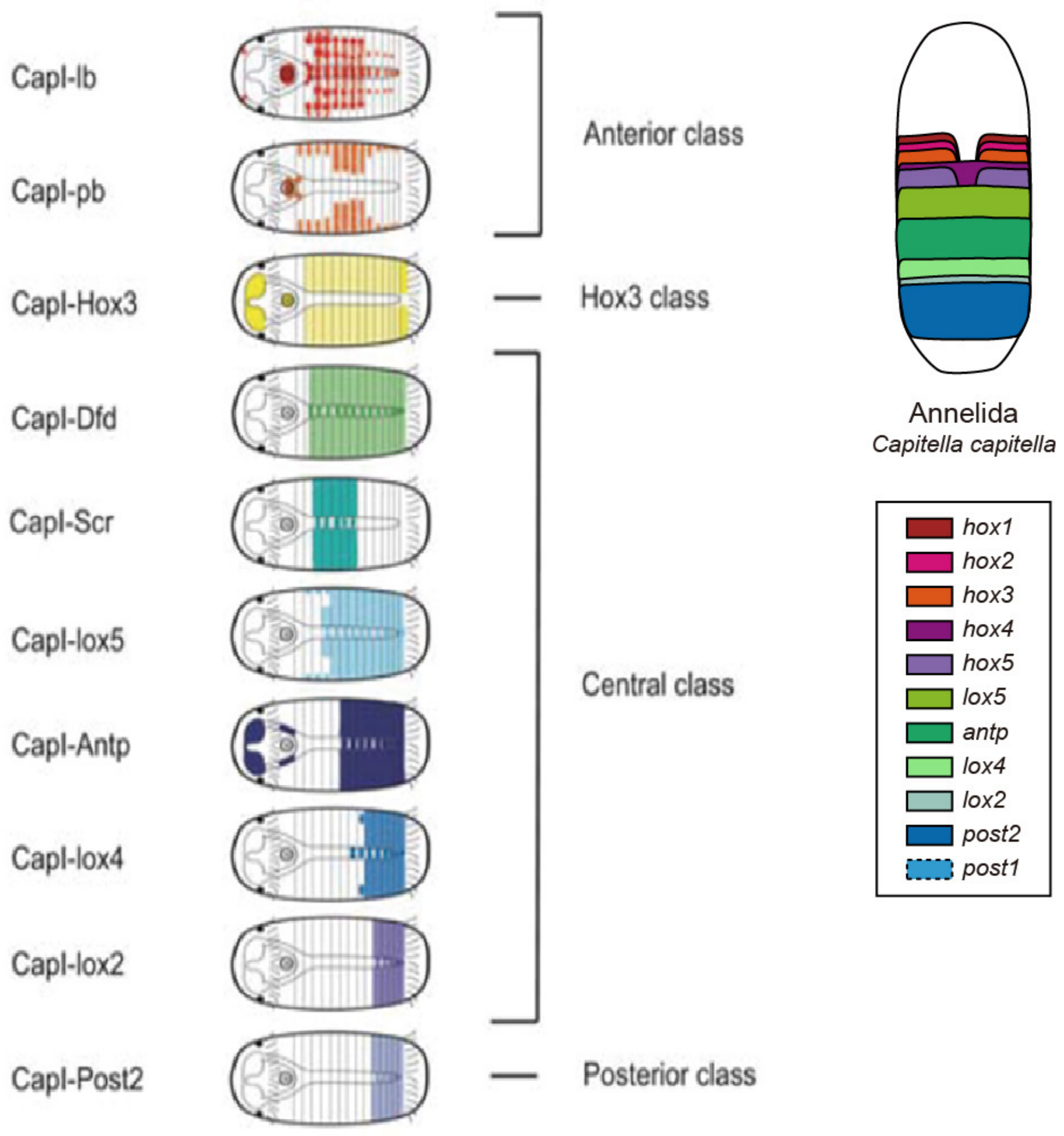


**Supplemental figure S12 The generally staggered** Hox **expression in the ventral tissues of the stage6-7 embryo of the annelid *Capitella teletAc.*** The original figure is published previously (Figure 14 in [4]) and reproduced here. A schematic diagram derived from these results is shown and also included in Fig. 7. Note that the colors we used to represent the Hox genes in the diagram are different from those used in the original article.

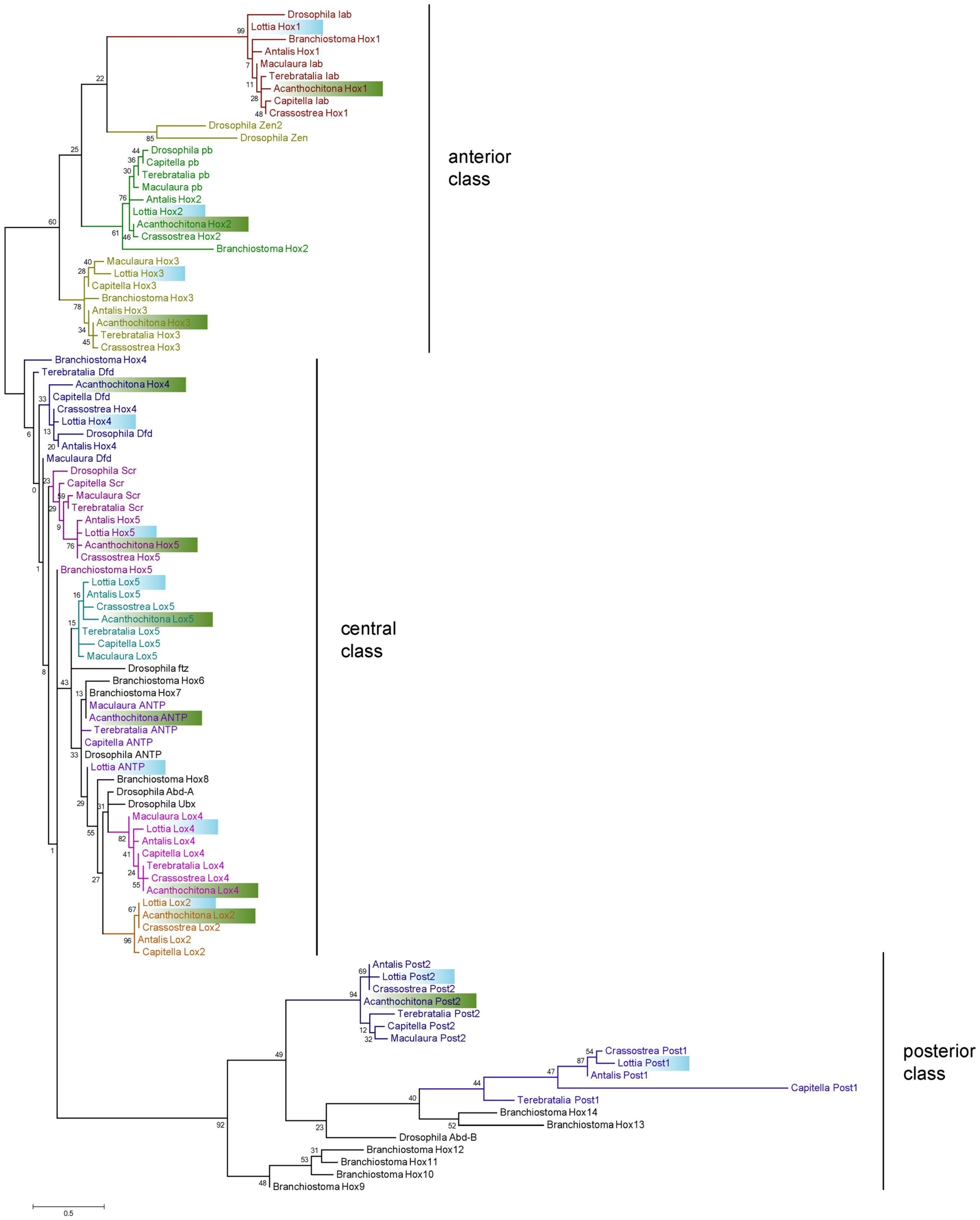


**Supplemental figure S12 ML tree of Hox genes.** The homeodomain of Hox genes from the two molluscan species we used in the study (indicated by blue and green boxes), as well as those from five other spiralians, an ecdysozoan and a deuterostome were used in the phylogenetic analysis (see details in supplemental table

S1). The orthologies of the genes are generally well discriminated despite relatively low resolutions for the central class genes. The phylogenetic analysis were performed using Mega6.0. The LG+G model was estimated to be the best fitting evolutionary model. The numbers at the nodes indicate bootstrap percentage from 1000 replicates.

```
Lottia_Hox1             PNSGRTNFTNKQLTELEKEFHFNKYLTRARRIEIAASLGLNETQVKIWFQNRRMKQKKRM
Acanthochitona_Hox1     PNMGRTNFTNKQLTELEKEFHYNKYLTRARRIEIAASLGLNETQVKIWFQNRRMKQKKRM
Crassostrea_Hox1        PNMGRTNFTNKQLTELEKEFHFNKYLTRARRIEIAAALGLNETQVKIWFQNRRMKQKKRL
Antalis_Hox1            ANGGRTNFTNKQLTELEKEFHFNKYLTRARRIEIAAALGLNETQVKIWFQNRRMKQKKRM
Capitella_lab           PNMGRTNFTNKQLTELEKEFHFNKYLTRARRIEIAASLGLNETQVKIWFQNRRMKQKKRL
Terebratalia_lab        PNMGRTNFSNKQLTELEKEFHFNKYLTRARRIEIAAALGLNETQVKIWFQNRRMKQKKRM
Maculaura_lab           PNTGRTNFTNKQLTELEKEFHFNKYLTRARRIEIAAALGLNETQVKIWFQNRRMKQKKRM

Lottia_Hox2             SRRLRTAYTNTQLLELEKEFHFNKYLCRPRRIEIAASLDLTERQVKVWFQNRRMKYKRQS
Acanthochitona_Hox2     TRRLRTAYTNTQLLELEKEFHFNKYLCRPRRIEIAASLDLTERQVKVWFQNRRMKYKRQS
Crassostrea_Hox2        TRRLRTAYTNTQLLELEKEFHFNKYLCRPRRIEIAASLDLTERQVKVWFQNRRMKYKRQT
Antalis_Hox2            GRRLRTAYTNTQLLELEKEFHFNKYLCRPRRIEIAASLELTERQVKVWFQNRRMKYKRQT
Capitella_pb            PRRLRTAYTNTQLLELEKEFHFNKYLCRPRRIEIAASLDLTERQVKVWFQNRRMKFKRQT
Terebratalia_pb         PRRLRTAYTNTQLLELEKEFHFNKYLCRPRRIEIAASLDLTERQVKVWFQNRRMKFKRQS
Maculaura_pb            PRRLRTAYTNSQLLELEKEFHFNKYLCRPRRIEIAASLDLTERQVKVWFQNRRMKYKRQS

Lottia_Hox3             AKRARTAYTSAQLVELEKEFHFNRYLCRPRRIEMAALLNLSERQIKIWFQNRRMKFKKDC
Acanthochitona_Hox3     SKRARTAYTSAQLVELEKEFHFNRYLCRPRRIEMAALLSLTERQIKIWFQNRRMKFKKEQ
Crassostrea_Hox3        TKRARTAYTSAQLVELEKEFHFNRYLCRPRRIEMAALLSLTERQIKIWFQNRRMKFKKEQ
Antalis_Hox3            SKRARTAYTSAQLVELEKEFHFNRYLCRPRRIEMAALLNLTERQIKIWFQNRRMKFKKEQ
Capitella_Hox3          SKRARTAYTSAQLVELEKEFHFNRYLCRPRRIEMAALLNLTERQIKIWFQNRRMKYKKDQ
Terebratalia_Hox3       SKRARTAYTSAQLVELEKEFHFNRYLCRPRRIEMAALLSLSERQIKIWFQNRRMKFKKEQ
Maculaura_Hox3          PKRSRTAYTSAQLVELEKEFHFNRYLCRPRRIEMAALLNLSERQIKIWFQNRRMKYKKDQ

Lottia_Hox4             SKRNRTAYTRHQVLELEKEFHFNRYLTRRRRIEIAHTLCLSERQIKIWFQNRRMKWKKEH
Acanthochitona_Hox4     TKRVRTAYTRHQVLELEKEFHFNRYLTRKRRIEIAHSLCLSERQIKIWFQNRRMKWKKEH
Crassostrea_Hox4        SKRNRTAYTRHQILELEKEFHFNRYLTRRRRIEIAHTLCLSERQIKIWFQNRRMKWKKEH
Antalis_Hox4            SKRNRTAYTRHQILELEKEFHFNRYLTRRRRIEIAHTLCLSERQIKIWFQNRRMKWKKEN
Capitella_Dfd           SKRTRTAYTRHQILELEKEFHFNRYLTRRRRIEIAHTLCLSERQIKIWFQNRRMKWKKEH
Terebratalia_Dfd        PKRSRTAYTRHQILELEKEFHFNRYLTRRRRIEIAHALCLTERQIKIWFQNRRMKWKKEH
Maculaura_Dfd           NKRTRTAYTRHQILELEKEFHFNRYLTRRRRIEIAHALCLTERQIKIWFQNRRMKWKKEH

Lottia_Hox5             SKRSRTSYTRHQTLELEKEFHYNKYLTRRRRIEIAHALNLTERQIKIWFQNRRMKWKKDH
Acanthochitona_Hox5     SKRSRTSYTRHQTLELEKEFHYNKYLTRRRRIEIAHSLNLTERQIKIWFQNRRMKWKKEH
Crassostrea_Hox5        SKRSRTSYTRHQTLELEKEFHYNKYLTRRRRIEIAHALNLTERQIKIWFQNRRMKWKKEH
Antalis_Hox5            TKRSRTSYTRHQTLELEKEFHYNKYLTRRRRIEIAHALNLTERQIKIWFQNRRMKWKKEH
Capitella_Scr           NKRTRTSYTRHQTLELEKEFHFNRYLTRRRRIEIAHSLNLTERQIKIWFQNRRMKWKKEH
Terebratalia_Scr        SKRTRTSYTRHQTLELEKEFHFNRYLTRRRRIEIAHALNLTERQIKIWFQNRRMKWKKEQ
Maculaura_Scr           SKRTRTSYTRYQTLELEKEFHFNRYLTRRRRIEIAHALNLTERQIKIWFQNRRMKWKKEQ

                                                                                 “Lox5 Parapeptide”
Lottia_Lox5             QKRTRQTYTRYQTLELEKEFHFNRYLTRRRRIEVAHMLGLTERQIKIWFQNRRMKWKKEN       NVSKLTGPD
Acanthochitona_Lox5     TKRTRQTYTRHQTLELEKEFHYNRYLTRRRRIEIAHMLGLTERQIKIWFQNRRMKWKKEN       NIQKLTGPD
Crassostrea_Lox5        QKRTRQTYTRYQTLELEKEFHFNRYLTRRRRIEIAHLLGLTERQIKIWFQNRRMKWKKDN       NIPKLTGPD
Antalis_Lox5            QKRTRQTYTRYQTLELEKEFHFNRYLTRRRRIEIAHMLGLTERQIKIWFQNRRMKWKKEN       NLEKLTGPD
Capitella_Lox5          QKRTRQTYTRYQTLELEKEFHYNRYLTRRRRIEIAHALQLTERQIKIWFQNRRMKYKKEN       NISKLTGPN
Terebratalia_Lox5       QKRTRQTYTRYQTLELEKEFHFNRYLTRRRRIEIAHALGLTERQIKIWFQNRRMKWKKEN       NLPKLTGPN
Maculaura_Lox5          QKRTRQTYTRYQTLELEKEFHFNKYLTRRRRIEIAHALGLTERQIKIWFQNRRMKWKKEN       NLQKLTGPN

Lottia_ANTP             RKRGRQTYTRYQTLELEKEFHFNRYLTRRRRIEIAHALCLTERQIKIWFQNRRMKWKKEA
Acanthochitona_ANTP     RKRGRQTYTRYQTLELEKEFHFNKYLTRRRRIEIAHALCLTERQIKIWFQNRRMKWKKEN
Capitella_ANTP          RKRGRQTYTRYQTLELEKEFHFNRYLTRRRRIEIAHALCLTERQIKIWFQNRRMKWKKEN
Terebratalia_ANTP       RKRGRQTYSRHQTLELEKEFHFNRYLTRRRRIEIAHALCLTERQIKIWFQNRRMKWKKEN
Maculaura_ANTP          RKRGRQTYTRYQTLELEKEFHFNKYLTRRRRIEIAHALCLTERQIKIWFQNRRMKWKKEN

                                                                                 “Ubd-A Parapeptide”
Lottia_Lox4             RRRGRQTYSRYQTLELEKEFQFNHYLTRKRRIEIAHTLCLTERQIKIWFQNRRMKMKKER       QAIKDINGDPK
Acanthochitona_Lox4     RRRGRQTYSRYQTLELEKEFQFNHYLTRKRRIEVAHALCLTERQIKIWFQNRRMKLKKER       QQIKELNDTYK
Crassostrea_Lox4        RRRGRQTYSRYQTLELEKEFQFNHYLTRKRRIEVAHSLCLTERQIKIWFQNRRMKLKKER       QAIKEINDNST
Antalis_Lox4            RRRGRQTYSRYQTLELEKEFQFNHYLTRKRRIEIAHSLCLTERQIKIWFQNRRMKLKKEK       QVIKGINDIST
Capitella_Lox4          RRRGRQTYSRYQTLELEKEFQFNHYLTRKRRIEIAHALCLTERQIKIWFQNRRMKLKKER       QQIKDLNGLDG
Terebratalia_Lox4       RRRGRQTYSRYQTLELEKEFQFNHYLTRKRRIEVAHALCLTERQIKIWFQNRRMKLKKER       QQIKELNDTYS
Maculaura_Lox4          RRRGRQTYSRYQTLELEKEFQFNHYLTRRRRIEIAHSLCLTERQIKIWFQNRRMKLKKER       QQIKELNDSTK

Lottia_Lox2             RRRGRQTYTRFQTLELEKEFKFNRYLTRRRRIELSHMLCLTERQIKIWFQNRRMKEKKEL       QAIKELNSQTR
Acanthochitona_Lox2     RRRGRQTYTRFQTLELEKEFKFNRYLTRRRRIELSHMLCLTERQIKIWFQNRRMKEKKEL       QAIKELNEQCR
Crassostrea_Lox2        RRRGRQTYTRFQTLELEKEFKFNRYLTRRRRIELSHMLCLTERQIKIWFQNRRMKEKKEL       QAIKELNEQSR
Antalis_Lox2            RRRGRQTYTRYQTLELEKEFKFNRYLTRRRRIELSHMLCLTERQIKIWFQNRRMKEKKEL       QAIKELNEQCR
Capitella_Lox2          RRRGRQTYTRYQTLELEKEFKFNRYLTRRRRIELSHMLCLTERQIKIWFQNRRMKEKKEI       QAIKELNEKEK

Lottia_Post2            GRKKRKPYTRYQTMVLENEFLSSSYITRQKRWEISCKLQLSERQVKVWFQNRRMKRKKLT
Acanthochitona_Post2    GRKKRKPYTRYQTMVLENEFLNNSYITRQKRWEISCKLQLTERQVKVWFQNRRMKRKKLN
Crassostrea_Post2       GRKKRKPYTRYQTMVLENEFLNSSYITRQKRWEISCKLQLSERQVKVWFQNRRMKRKKLN
Antalis_Post2           GRKKRKPYTRYQTMVLENEFLNSSYITRQKRWEISCKLQLSERQVKVWFQNRRMKRKKLN
Capitella_Post2         QRKKRKPYTRYQTMVLENEFINNSYITRQKRWEISCKLHLSERQVKVWFQNRRMKRKKLN
Terebratalia_Post2      SRKKRKPYTRYQNMLLENEFIANSYITRQKRWEISCKLQLTERQVKVWFQNRRMKRKKLT
Maculaura_Post2         TRKKRKPYTRYQTMVLENEFMTNSYITRQKRWEISCKLHLTERQVKVWFQNRRMKRKKLN

Lottia_Post1            LRKRRRPYSKFQIAELEREYNGSTYVSKSRRWELSQLINLSERQIKIWFQNRRIKAKKII
Crassostrea_Post1       LRKRRRPYSKFQIAELEREYNNSTYISKSRRWELSQLINLSERQIKIWFQNRRIKAKKVS
Antalis_Post1           LRKRRRPYSKFQIAELEREYANSTYISKSRRWELSQLINLSERQIKIWFQNRRIKAKKIQ
Capitella_Post1         PKKKRKPYSKPQVSALENEYSTSTYITKARRKEVARELDLTERQIKIWYQNRRIKEKKIA
Terebratalia_Post1      MRKRRKPYSKQQINELEREYVKTTYISKPKRWELAQRLNLSERQVKIWFQNRRMKEKKMR
```

**Supplemental figure S13 Alignment of homeodomains from the two mollusks in this study as well as other spiralians.** The diagnostic amino acid residues are highlighted according to previous reports [5-7]**.** The parapeptides from the three central class Hox genes (Lox5, Lox4 and Lox2) are also shown.

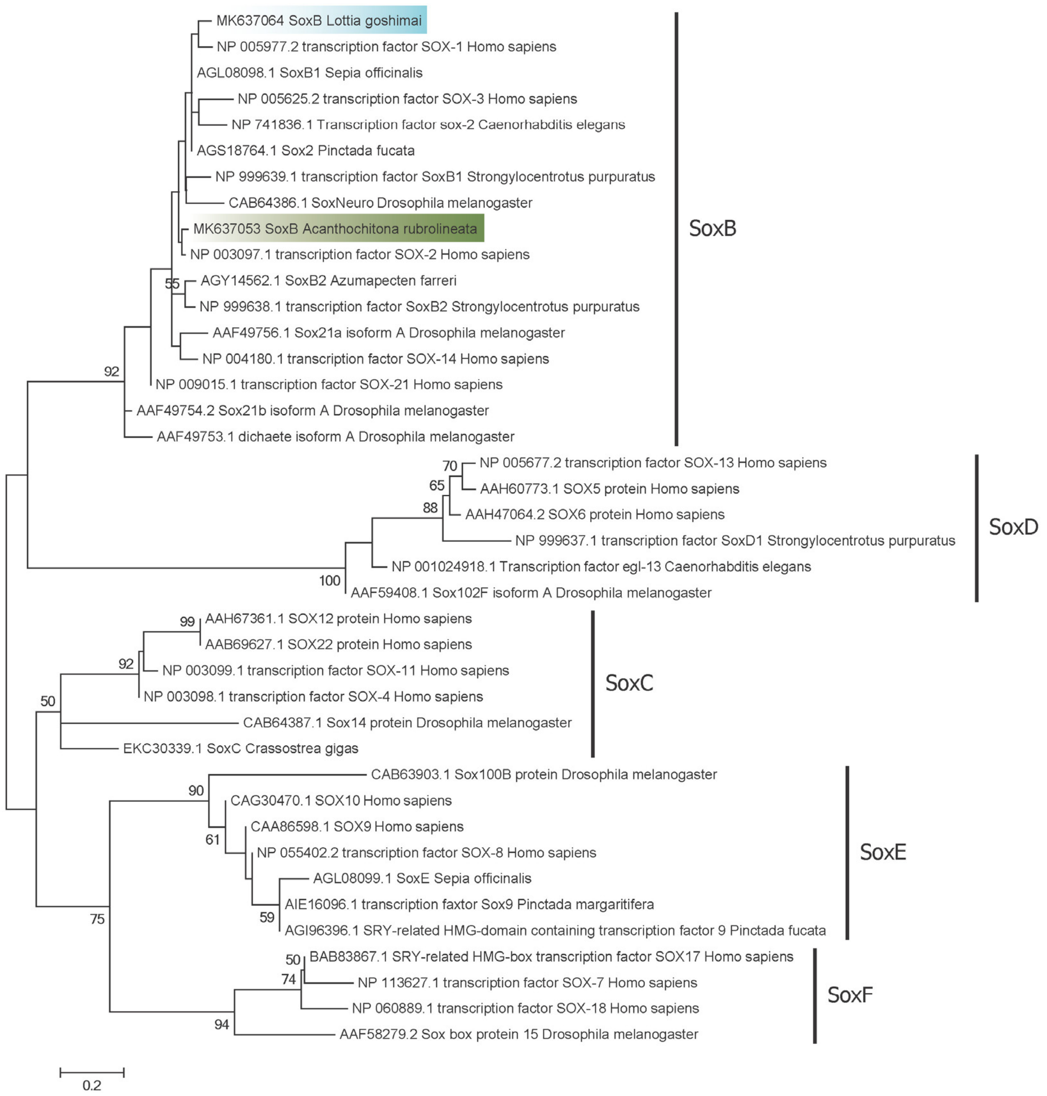


**Supplemental figure S14 ML tree of SoxB.** The HMG domain of Sox genes from major SOX sub-families were used in the phylogenetic analysis. The phylogenetic analysis were performed using Mega6.0. The LG+G model was estimated to be the best fitting evolutionary model. The numbers at the nodes indicate bootstrap percentage from 1000 replicates.

**Supplemental Table S1. Hox genes used in supplemental figures S12 and S13.**

| Species | Gene | Accession | Abbreviation |
|---|---|---|---|
| *Branchiostoma floridae* | Hox1 | BAA78620.2 | Branchiostoma Hox1 |
| *Branchiostoma floridae* | Hox2 | BAA78621.1 | Branchiostoma Hox2 |
| *Branchiostoma floridae* | Hox3 | X68045.1 | Branchiostoma Hox3 |
| *Branchiostoma floridae* | Hox4 | BAA78622.1 | Branchiostoma Hox4 |
| *Branchiostoma floridae* | Hox5 | CAA84517.1 | Branchiostoma Hox5 |
| *Branchiostoma floridae* | Hox6 | CAA84518.1 | Branchiostoma Hox6 |
| *Branchiostoma floridae* | Hox7 | CAA84519.1 | Branchiostoma Hox7 |
| *Branchiostoma floridae* | Hox8 | CAA84520.1 | Branchiostoma Hox8 |
| *Branchiostoma floridae* | Hox9 | CAA84521.1 | Branchiostoma Hox9 |
| *Branchiostoma floridae* | Hox10 | CAA84522.1 | Branchiostoma Hox10 |
| *Branchiostoma floridae* | Hox11 | AAF81909.1 | Branchiostoma Hox11 |
| *Branchiostoma floridae* | Hox12 | AAF81903.1 | Branchiostoma Hox12 |
| *Branchiostoma floridae* | Hox13 | AAF81904.1 | Branchiostoma Hox13 |
| *Branchiostoma floridae* | Hox14 | AAF81905.1 | Branchiostoma Hox14 |
| *Drosophila melanogaster* | lab | CAB57787.1 | Drosophila lab |
| *Drosophila melanogaster* | pb | CAA45271.1 | Drosophila pb |
| *Drosophila melanogaster* | Zen | AAF54087.1 | Drosophila Zen |
| *Drosophila melanogaster* | Zen2 | P09090.2 | Drosophila Zen2 |
| *Drosophila melanogaster* | Dfd | P07548.2 | Drosophila Dfd |
| *Drosophila melanogaster* | Scr | NP 524248.2 | Drosophila Scr |
| *Drosophila melanogaster* | ftz | NP 477498.1 | Drosophila ftz |
| *Drosophila melanogaster* | ANTP | CAA27417.1 | Drosophila ANTP |
| *Drosophila melanogaster* | Ubx | CAA29194.1 | Drosophila Ubx |
| *Drosophila melanogaster* | Abd-A | P29555.2 | Drosophila Abd-A |
| *Drosophila melanogaster* | Abd-B | CAB57859.1 | Drosophila Abd-B |
| *Capitella teleta* | lab | EU196537.1 | Capitella lab |
| *Capitella teleta* | pb | EU196538.1 | Capitella pb |
| *Capitella teleta* | Hox3 | EU196539.1 | Capitella Hox3 |
| *Capitella teleta* | Dfd | EU196540.1 | Capitella Dfd |
| *Capitella teleta* | Scr | EU196541.1 | Capitella Scr |
| *Capitella teleta* | Lox5 | EU196542.1 | Capitella Lox5 |
| *Capitella teleta* | ANTP | EU196547.1 | Capitella ANTP |
| *Capitella teleta* | Lox4 | EU196543.1 | Capitella Lox4 |
| *Capitella teleta* | Lox2 | EU196544.1 | Capitella Lox2 |
| *Capitella teleta* | Post2 | EU196545.1 | Capitella Post2 |
| *Capitella teleta* | Post1 | EU196546.1 | Capitella Post1 |
| *Maculaura alaskensis* | lab | KP762174.1 | Maculaura lab |
| *Maculaura alaskensis* | pb | KP762176.1 | Maculaura pb |
| *Maculaura alaskensis* | Hox3 | KP762173.1 | Maculaura Hox3 |
| *Maculaura alaskensis* | Dfd | KP762180.1 | Maculaura Dfd |

| | | | |
|---|---|---|---|
| *Maculaura alaskensis* | Scr | KP762177.1 | Maculaura Scr |
| *Maculaura alaskensis* | Lox5 | KP762179.1 | Maculaura Lox5 |
| *Maculaura alaskensis* | ANTP | KP762171.1 | Maculaura ANTP |
| *Maculaura alaskensis* | Lox4 | KP762175.1 | Maculaura Lox4 |
| *Maculaura alaskensis* | Post2 | KP762178.1 | Maculaura Post2 |
| *Terebratalia transversa* | lab | KX372761.1 | Terebratalia lab |
| *Terebratalia transversa* | pb | KX372762.1 | Terebratalia pb |
| *Terebratalia transversa* | Hox3 | KX372763.1 | Terebratalia Hox3 |
| *Terebratalia transversa* | Dfd | KX372764.1 | Terebratalia Dfd |
| *Terebratalia transversa* | Scr | KX372765.1 | Terebratalia Scr |
| *Terebratalia transversa* | Lox5 | KX372766.1 | Terebratalia Lox5 |
| *Terebratalia transversa* | ANTP | KX372771.1 | Terebratalia ANTP |
| *Terebratalia transversa* | Lox4 | KX372767.1 | Terebratalia Lox4 |
| *Terebratalia transversa* | Post2 | KX372768.1 | Terebratalia Post2 |
| *Terebratalia transversa* | Post1 | KX372772.1 | Terebratalia Post1 |
| *Antalis entalis* | Hox1 | KX365088.1 | Antalis Hox1 |
| *Antalis entalis* | Hox2 | KX365089.1 | Antalis Hox2 |
| *Antalis entalis* | Hox3 | KX365090.1 | Antalis Hox3 |
| *Antalis entalis* | Hox4 | KX365091.1 | Antalis Hox4 |
| *Antalis entalis* | Hox5 | KX365092.1 | Antalis Hox5 |
| *Antalis entalis* | Lox5 | KX365093.1 | Antalis Lox5 |
| *Antalis entalis* | Lox4 | KX365095.1 | Antalis Lox4 |
| *Antalis entalis* | Lox2 | KX365094.1 | Antalis Lox2 |
| *Antalis entalis* | Post2 | KX365097.1 | Antalis Post2 |
| *Antalis entalis* | Post1 | KX365096.1 | Antalis Post1 |
| *Crassostrea gigas** | Hox1 | CGI 10024083 | Crassostrea Hox1 |
| *Crassostrea gigas** | Hox2 | CGI 10024086 | Crassostrea Hox2 |
| *Crassostrea gigas** | Hox3 | CGI 10024087 | Crassostrea Hox3 |
| *Crassostrea gigas** | Hox4 | CGI 10024091 | Crassostrea Hox4 |
| *Crassostrea gigas* | Hox5 | XP 011424787.1 | Crassostrea Hox5 |
| *Crassostrea gigas** | Lox5 | CGI 10026565 | Crassostrea Lox5 |
| *Crassostrea gigas** | Lox4 | CGI 10026562 | Crassostrea Lox4 |
| *Crassostrea gigas** | Lox2 | CGI 10018592 | Crassostrea Lox2 |
| *Crassostrea gigas** | Post2 | CGI 10027388 | Crassostrea Post2 |
| *Crassostrea gigas** | Post1 | CGI 10027385 | Crassostrea Post1 |
| *Lottia goshimai* | Hox1 | MK637065 | Lottia Hox1 |
| *Lottia goshimai* | Hox2 | MK637066 | Lottia Hox2 |
| *Lottia goshimai* | Hox3 | MK637067 | Lottia Hox3 |
| *Lottia goshimai* | Hox4 | MK637068 | Lottia Hox4 |
| *Lottia goshimai* | Hox5 | MK637069 | Lottia Hox5 |
| *Lottia goshimai* | Lox5 | MK637070 | Lottia Lox5 |
| *Lottia goshimai* | ANTP | MK637071 | Lottia ANTP |
| *Lottia goshimai* | Lox4 | MK637072 | Lottia Lox4 |
| *Lottia goshimai* | Lox2 | MK637073 | Lottia Lox2 |

| | | | |
|---|---|---|---|
| *Lottia goshimai* | Post2 | MK637074 | Lottia Post2 |
| *Lottia goshimai* | Post1 | MK637075 | Lottia Post1 |
| *Acanthochitona rubrolineata* | Hox1 | MK637054 | Acanthochitona Hox1 |
| *Acanthochitona rubrolineata* | Hox2 | MK637055 | Acanthochitona Hox2 |
| *Acanthochitona rubrolineata* | Hox3 | MK637056 | Acanthochitona Hox3 |
| *Acanthochitona rubrolineata* | Hox4 | MK637057 | Acanthochitona Hox4 |
| *Acanthochitona rubrolineata* | Hox5 | MK637058 | Acanthochitona Hox5 |
| *Acanthochitona rubrolineata* | Lox5 | MK637059 | Acanthochitona Lox5 |
| *Acanthochitona rubrolineata* | ANTP | MK637060 | Acanthochitona ANTP |
| *Acanthochitona rubrolineata* | Lox4 | MK637061 | Acanthochitona Lox4 |
| *Acanthochitona rubrolineata* | Lox2 | MK637062 | Acanthochitona Lox2 |
| *Acanthochitona rubrolineata* | Post2 | MK637063 | Acanthochitona Post2 |

*: These sequences were retrieved from OysterDB (http://www.oysterdb.com/).

### Supplemental Table S2. Primers used in this study.

| primer | sequence (5'-3') | primer | sequence (5'-3') |
|---|---|---|---|
| LgHox1-F | TCGGCTAATGGAACCTGTATG | LgHox1-R | GTGTATCATGACACATGGCTAA |
| LgHox2-F | TGAACGAGGAGGGTGAATGTG | LgHox2-R | AGAACGTCCTGACTGAGATTGG |
| LgHox3-F | TTACGCTTCTTACGGCACGAT | LgHox3-R | TCAGAACCACCTTTCAGTCTACAG |
| LgHox4-F | CCGTCAGAGGAGTATTCACAATCT | LgHox4-R | CTGTCGGACTCATATCGTCATCA |
| LgHox5-F | GCCGCTTATTATCCACACAAACA | LgHox5-R | TCACCTTCTCTAGGACTAGGACAT |
| LgLox5-F | TATGCGATGTAGCGGTTATAGTGA | LgLox5-R | TGTTAGCGAAGAACTGCCAATATC |
| LgAntp-F | ATGGACCTGATTATGCCAGTGTT | LgAntp-R | TTTTTCATCCTCTAAATCGTCACT |
| LgLox4-F | TCCACAGGACATTACTATCCACAA | LgLox4-R | GGTTTCAACTCGAATTCTCGTTTG |
| LgLox2-F | CACTGGAATAACCGAAGATTGTCA | LgLox2-R | GGACGTGTTTGGGAATTAAGTTCT |
| LgPost2-F | GGCGAATAACATTCTGGCTCATCC | LgPost2-R | TCCACTGCCGTACAATGTCTAATATG |
| LgPost1-F | CAGCAGTGTATCCGTTCCAATC | LgPost1-R | TGACGCTCTGATAGGTTGATAAGT |
| ArHox1-F | TTCCGAACTGACTGTGTCTTGTG | ArHox1-R | TGTGCCTCTTTCATCCGTTTCTT |
| ArHox2-F | GCGTGGAGAGGACTAACTGTG | ArHox2-R | GCCGTTGGAGGTTGGATACATAT |
| ArHox3-F | ACGAACCAGTTCAAACTTTCCCT | ArHox3-R | TTTGTCTGAGTCGCCTTTACCTTT |
| ArHox4-F | CGCCATACGCAGATCCTAAGTT | ArHox4-R | ACCAAGGTAACCGTCCGAAGA |
| ArHox5-F | GCCATCCGAATTACAGTCATCCT | ArHox5-R | GGCTATGTGTGATAGTTTGTGCTC |
| ArLox5-F | GCAACTCGTACTCAACTCACTCC | ArLox5-R | CCATGCTGAACACTTCTAGAGTCT |
| ArAntp-F | ACGGAGTCTACTTTCTACCACCAT | ArAntp-R | CGGGCGAGTCGAGTAATTTGTT |
| ArLox4-F | GAGTCCGCACGGCGAGAA | ArLox4-R | TCGTCCACATCCTCCTTAATATCG |
| ArLox2-F | CCTACCCACTCCTTCGCCATA | ArLox2-R | CAACTGAACTCCTGTATCGTCCTT |
| ArPost2-F | CTGTTGTCACTTCATCGAGCATTC | APost2-R | AGAGCCATTATTCATGGTCACTGT |
| LgSoxb-F | TTGCGACCGATCCTAGAGTGA | LgSoxb-R | TCCGTCTGTGGTGGCTTGT |
| ArSoxb-F | TCAGTTCAGAGAATGGAGACTACAGAT | ArSoxb-R | CGGCATAGAGTAAGGCGACATTG |

### Supplemental references


1. Fröbius AC, Funch P: **Rotiferan Hox genes give new insights into the evolution of metazoan bodyplans**. *Nature Communications* 2017, **8**(1).
2. Hiebert LS, Maslakova SA: **Expression of Hox, Cdx, and Six3/6 genes in the hoplonemertean *Pantinonemertes californiensis* offers insight into the evolution of maximally indirect development in the phylum Nemertea.** *EvoDevo* 2015, **6**(1):1-15.
3. Schiemann SM, Martín-Durán JM, Børve A, Vellutini BC, Passamaneck YJ, Hejnol A: **Clustered brachiopod Hox genes are not expressed collinearly and are associated with lophotrochozoan novelties**. *Proceedings of the National Academy of Sciences* 2017, **114**(10):E1913-E1922.
4. Fröbius AC, Matus DQ, Seaver EC: **Genomic organization and expression demonstrate spatial and temporal Hox gene colinearity in the lophotrochozoan *Capitella sp. I***. *PLoS ONE* 2008, **3**(12).
5. Balavoine G, De Rosa R, Adoutte A: **Hox clusters and bilaterian phylogeny**. *Molecular Phylogenetics and Evolution* 2002, **24**(3):366-373.
6. Matus DQ, Halanych KM, Martindale MQ: **The Hox gene complement of a pelagic chaetognath, *Flaccisagitta enflata*.** *Integrative and Comparative Biology* 2007, **47**(6):854-864.
7. Pérez-Parallé ML, Pazos AJ, Mesías-Gansbiller C, Sánchez JL: **Hox, ParaHox, Ehgbox, and NK Genes in Bivalve Molluscs: Evolutionary Implications**. *Journal of Shellfish Research* 2016, **35**(1):179-190.
